# GTX-GUT: A Standardized Metagenomic Workflow for Gut Microbiome Profiling and Clinical Associations

**DOI:** 10.64898/2026.08.12.744496

**Authors:** Rodrigo Lima Andrade, Tayná da Silva Fiúza, José Eduardo Kroll, Pedro Victor Barbosa Araújo, Daniel Henrique Ferreira Gomes, Leonardo Varuzza, Gustavo Antonio de Souza, Pitágoras de A. Alves Sobrinho, Sandro José de Souza

## Abstract

The human gut microbiome plays a central role in host physiology and disease, yet metagenomic analysis pipelines remain fragmented across sample preparation, taxonomic classification, and clinical interpretation stages, complicating reproducibility and translational use. Here we present GTX-GUT, a fully automated, containerized Snakemake pipeline for 16S rRNA gut microbiome profiling that integrates quality control, taxonomic classification (QIIME2/DADA2 against Greengenes 13.8), diversity and compositional metrics benchmarked against a curated healthy reference population, enterotype classification, a clinical association module spanning 11 disease categories, and automated natural-language report generation. We validated the pipeline using the ZymoBIOMICS mock community, showing that BBDuk preprocessing substantially reduced genus-level quantification error (Mean Absolute Error reduced from 7.34 to 1.58 percentage points; Pearson’s *r* improved from 0.576 to 0.833). Application to a human sample from a patient with type 2 Diabetes Mellitus recovered a dysbiotic signature consistent with the literature, including reduced Firmicutes abundance, elevated Bacteroidetes and Proteobacteria, and a predominance of clinical associations within metabolic and gastrointestinal categories. These results demonstrate that GTX-GUT provides a reproducible, end-to-end framework linking raw sequencing data to clinically interpretable output, with direct applicability to research and translational microbiome studies.

## Introduction

The human gut microbiome has emerged as a central focus in biomedical research due to its profound influence on host physiology, metabolism, immunity, and disease susceptibility. Comprising trillions of microorganisms—including bacteria, archaea, viruses, and fungi—the gut microbiome represents a highly complex ecosystem whose composition and functional dynamics are tightly linked to health and disease states. Understanding this intricate microbial community requires robust analytical approaches capable of capturing both taxonomic diversity and functional potential.

Metagenomics, the culture-independent study of microbial genomes directly from environmental samples, has revolutionized microbiome research. By sequencing and analyzing the collective genetic material of gut microorganisms, metagenomics enables comprehensive profiling of microbial communities, including rare or uncultivable taxa. This approach not only provides insights into microbial composition but also allows functional characterization through gene and pathway analysis, thereby bridging the gap between microbial presence and their biological roles (1).

Despite its transformative potential, metagenomic analysis of the gut microbiome presents significant challenges (2, 3). These include the complexity of sample preparation, the vast amount of sequencing data generated, and the need for sophisticated computational pipelines to ensure accurate taxonomic classification, functional annotation, and statistical interpretation. Moreover, variability in methodological choices—such as DNA extraction protocols, sequencing platforms, and bioinformatic tools—can introduce biases that complicate reproducibility and comparability across studies.

To address these challenges, standardized pipelines for gut microbiome analysis are essential. Such pipelines integrate multiple stages, including quality control of raw sequencing data, assembly or mapping strategies, taxonomic profiling, functional annotation, and statistical modeling. By harmonizing these steps, researchers can achieve more reliable and reproducible results, facilitating cross-study comparisons and enabling translational applications in clinical and nutritional sciences.

This paper presents GTX-GUT, a comprehensive, fully automated pipeline for the analysis of the gut microbiome based on 16S rRNA metagenomics. The proposed workflow emphasizes methodological rigor, reproducibility, and adaptability, providing a framework that can be applied to diverse datasets and directly linked to clinical interpretation. Ultimately, the development and dissemination of standardized pipelines will accelerate discoveries in microbiome science and foster their integration into precision medicine and public health.

## Methods

### Pipeline Architecture and Workflow Management

The GTX-GUT pipeline is implemented as a fully automated Snakemake workflow (4), version *≥* 8.27, which orchestrates all analytical stages—from raw sequencing read processing to the generation of a clinical report—without manual intervention between steps. Each stage is executed inside dedicated Singularity-CE containers, version 3.11 (5), ensuring reproducibility and portability of the software environment across computing infrastructures. Fig. 1 summarizes the overall architecture of the pipeline, comprising quality-controlled preprocessing, QIIME2-based taxonomic classification, abundance-based statistical evaluation, and automated report generation modules, including the clinical association and interpretive text-generation components.

**Fig 1.**
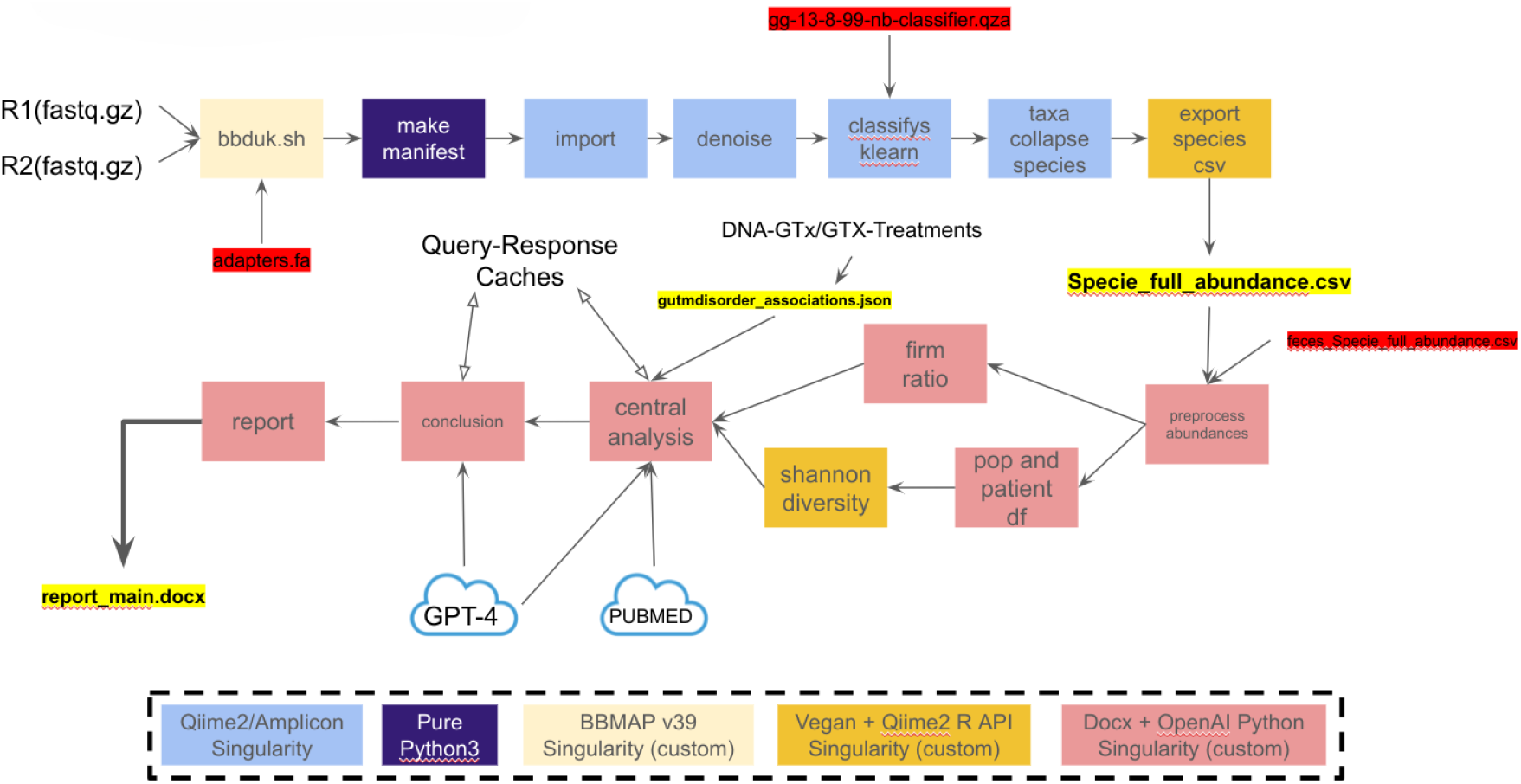
Overview of the GTX-GUT pipeline architecture. Raw single-end or paired-end reads are quality-filtered with BBDuk, imported and denoised in QIIME2, and classified against the Greengenes 13.8 naive Bayes classifier. Species-level abundance tables are preprocessed and used to compute the Firmicutes/Bacteroidetes ratio, Shannon diversity, and population/patient comparisons in the central analysis module, which queries a curated gut-disorder association database and large language model (GPT-4) and PubMed-backed caches to generate an interpretive conclusion and a consolidated .docx clinical report.

### Sequencing Data and Preprocessing

The pipeline accepts 16S rRNA gene amplicon sequencing reads in FASTQ format, supporting both single-end and paired-end sequencing layouts. Because taxonomic classification relies on a reference classifier trained on the full-length 16S rRNA gene (see below), input samples are not restricted to a specific hypervariable region or primer pair. Prior to taxonomic classification, raw reads are quality-filtered and adapter-trimmed using BBDuk, part of the BBTools suite (6), version 39. This step removes low-quality bases, sequencing adapters, and contaminant reads, and its impact on downstream classification accuracy was explicitly quantified as part of the functional validation protocol described below.

### Taxonomic Classification

Taxonomic assignment is performed using QIIME 2 (7), version 2023.9, with amplicon sequence variants (ASVs) inferred through the DADA2 algorithm (8), which corrects sequencing errors and resolves single-nucleotide differences between sequences. ASVs are classified against the Greengenes reference database, version 13.8 (9), at 99% sequence similarity, using a naive Bayes classifier trained on the full-length 16S rRNA gene, which allows classification of amplicons generated with different hypervariable regions and primer pairs. Taxonomic profiles are aggregated at the phylum and genus levels for downstream diversity and compositional analyses.

### Diversity, Compositional, and Reference-Based Metrics

For each sample, the pipeline computes the Shannon diversity index (10) and the Firmicutes/Bacteroidetes (F/B) ratio, two widely used indicators of gut microbial ecosystem health. Both metrics are evaluated against fixed, literature-informed reference intervals rather than the empirical distribution of the reference population, allowing consistent interpretation across samples independent of cohort composition. Phylum-level abundances are additionally classified into three statistical zones—Very Low, Normal, and Very High—based on their position relative to the interquartile range of a curated reference population (described below), enabling automated flagging of taxa that deviate substantially from typical healthy-gut composition.

### Reference Population

A reference population of healthy gut microbiome profiles was constructed from publicly available 16S rRNA sequencing data (BioProject PRJEB53463, USDA Western Human Nutrition Research Center), generated on the Illumina MiSeq platform targeting the V4–V5 hypervariable regions (300 bp paired-end reads), to contextualize sample-level results. Of the 530 sequencing runs available in the BioProject, runs with near-zero sequencing depth were excluded (n = 501), followed by deduplication of repeated sampling from the same host individual (n = 401) and further quality-control filtering intrinsic to the pipeline (n = 372), yielding a final reference population of 350 healthy individuals. Distributions of Shannon diversity and F/B ratio within the resulting reference population were tested for unimodality using Hartigan’s dip test (11), supporting their use as a baseline for the classification framework described above.

### Enterotype Classification

Samples are classified into one of three enterotypes—*Bacteroides*-, *Prevotella*-, or *Ruminococcus*-dominated—using a simplified dominant-genus approach, in which the enterotype is assigned according to the most abundant of the three defining genera (12). This approach was adopted in place of the original Jensen–Shannon distance and partitioning-around-medoids (PAM) clustering method to allow rapid, single-sample classification without requiring a full reference cohort at runtime.

### Clinical Association Module

Taxa identified in each sample are cross-referenced against a curated database of taxon– condition associations spanning 11 clinical categories (e.g., gastrointestinal, metabolic and endocrine, psychiatric and neuropsychiatric, and neoplastic conditions), compiled from the peer-reviewed literature and informed by the gutMDisorder database (13). For each association, the pipeline reports the direction of the relationship (positive or negative), the statistical positioning of the taxon relative to the reference population, and a quantitative summary of the number of alterations detected per category.

### Automated Report Generation

At the conclusion of processing, the pipeline compiles all quantitative and statistical outputs into a structured clinical report in .docx format. A textual synthesis of the findings is generated using the GPT-4o-mini model accessed via the OpenAI API (14), which produces an interpretive conclusion summarizing diversity metrics, phylum-level alterations, enterotype classification, and clinical associations in natural language.

### Functional Validation Protocol

To assess the accuracy of the preprocessing and taxonomic classification steps, the pipeline was validated using the ZymoBIOMICS Microbial Community Standard (Catalog No. D6300; (15)), a mock community with a manufacturer-defined nominal composition of eight bacterial genera. Sequencing data from the mock community were processed through the full GTX-GUT pipeline both with and without the BBDuk preprocessing step, and the resulting taxonomic profiles were compared against the nominal composition using absolute error pergenus, Mean Absolute Error (MAE), Root Mean Square Error (RMSE), and Pearson’s correlation coefficient, following recommended benchmarking practices for omics computational tools (16).

### Computational Environment and Reproducibility

All analyses were executed within Singularity-CE containers (version 3.11) (5) orchestrated by Snakemake (version *≥* 8.27) (4), using Python 3.11 (with matplotlib 3.10 and the OpenAI SDK) and R with Bioconductor for statistical analysis and visualization. This containerized, workflow-managed design ensures that the entire pipeline—from raw sequencing reads to the final clinical report—can be executed deterministically on any compatible computing environment, addressing the reproducibility challenges that motivate the development of standardized metagenomic pipelines (2, 17).

## Availability

The version of GTX-GUT used in this report is available for non-commercial usage upon request.

## Results

### Functional Validation of the Pipeline

Functional validation of the pipeline was performed using the ZymoBIOMICS Microbial Community Standard (mock community), whose taxonomic profile is well-characterized and documented by the manufacturer. This approach allows verification of whether the preprocessing and taxonomic classification steps yield results consistent with the expected composition (16).

#### Impact of Preprocessing on Taxonomic Classification

For validation, we employed the ZymoBIOMICS Microbial Community Standard (D6300), a microbial community reference with a nominal composition defined and documented by Zymo Research (15). This standard consists of eight bacterial genera in predetermined proportions. Table 1 presents the expected relative abundances (nominal composition of the sample) compared with the results obtained using the GTX-GUT pipeline, both with and without the filtering step performed by BBDuk version 39.

**Table 1.** Expected relative abundances (nominal composition declared by the manufacturer, ZymoBIOMICS D6300) and observed abundances, with and without BBDuk preprocessing, for the eight bacterial genera comprising the mock community standard.

| Taxon | Expected (%) | Filtered (%) | Unfiltered (%) |
| --- | --- | --- | --- |
| *Bacillus* | 17.4 | 18.00 | 44.76 |
| *Listeria* | 14.1 | 14.06 | 8.34 |
| *Staphylococcus* | 15.5 | 15.69 | 15.45 |
| *Lactobacillus* | 18.4 | 12.08 | 6.12 |
| *Enterococcus* | 9.9 | 11.75 | 7.44 |
| *Salmonella* | 10.4 | 11.69 | 6.99 |
| *Escherichia (E. coli)* | 10.1 | 11.14 | 6.89 |
| *Pseudomonas* | 4.2 | 5.51 | 0.00 |

Fig. 2 illustrates the absolute error between observed and expected abundances for each genus, comparing results obtained with and without the preprocessing step. Without filtering, the abundance of *Bacillus* showed a pronounced distortion, reaching approximately 44.76% compared to the expected 17.4%—an absolute error of 27.36 percentage points. After applying BBDuk, this abundance was reduced to 18.00%, resulting in an absolute error of only 0.60 percentage points. A similar pattern of error reduction was observed across most genera, with notable improvements for *Listeria* (error reduced from 5.76 to 0.04 percentage points) and *Salmonella* (from 3.41 to 1.29 percentage points).

**Fig 2.**
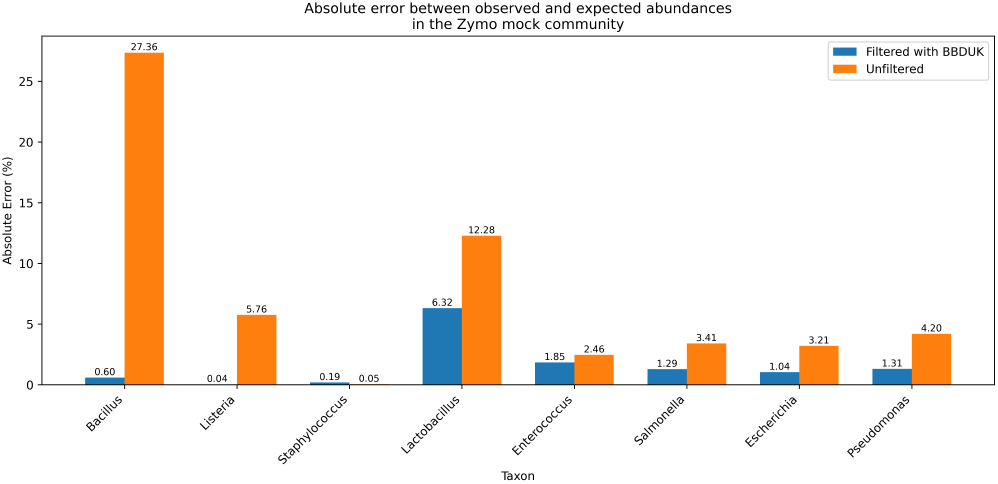
Absolute error (percentage points) between observed and expected genus-level abundances in the ZymoBIOMICS mock community, comparing results with and without BBDuk preprocessing.

To summarize the overall impact of preprocessing on taxonomic classification accuracy, three concordance metrics were calculated between the observed and nominal compositions: Mean Absolute Error (MAE), Root Mean Square Error (RMSE), and Pearson’s correlation coefficient. Table 2 presents these results, highlighting the substantial improvement in accuracy achieved through the preprocessing step.

**Table 2.** Concordance metrics between observed and manufacturer-declared abundances for the ZymoBIOMICS D6300 sample, calculated with and without BBDuk preprocessing.

| Method | MAE | RMSE | Pearson's *r* |
| --- | --- | --- | --- |
| Filtered (BBDuk) | 1.58 | 2.46 | 0.833 |
| Unfiltered | 7.34 | 11.06 | 0.576 |

### Application to a Human Sample

The following results were obtained using the GTX-GUT pipeline for the human sample SRR33578315, derived from a patient diagnosed with type 2 Diabetes Mellitus. The dataset was processed through the pipeline in a fully automated manner, ensuring consis-tency and reproducibility across all analytical steps.

The Shannon diversity index (10) calculated for the sample was 2.571. This value places the sample within the medium-to-low diversity range (2 *≤ H*, *<* 3), which is below the typical range observed in healthy reference populations. Such a reduction may reflect the characteristic microbial profile of patients with type 2 Diabetes Mellitus, a condition frequently associated with decreased alpha diversity in the gut microbiome.

The Firmicutes/Bacteroidetes (F/B) ratio calculated for the sample was 0.562, which falls within the clinically balanced range (F/B *<* 1.5). Although technically within the equilibrium threshold, this markedly reduced value compared to healthy reference populations reflects the dominance of *Bacteroidetes* observed in the sample. Such a pattern is consistent with microbial alterations frequently reported in patients with type 2 Diabetes Mellitus (18, 19).

Fig. 3 shows the relative abundances of the major bacterial phyla identified in the sample, along with their statistical classifications compared to the healthy reference population. The phylum *Firmicutes* exhibited an abundance of 31.86% (classified as *Very Low*), well below its typical predominance in the healthy gut microbiota, where the median reference value is approximately 85%. In contrast, *Bacteroidetes* displayed a markedly elevated abundance of 56.65% (*Very High*), far exceeding reference values.

**Fig 3.**
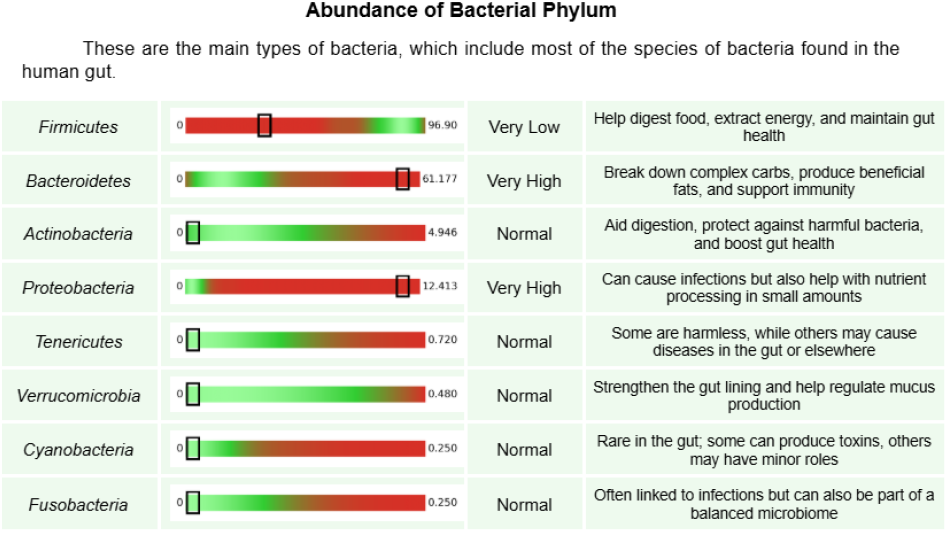
Relative abundance of major bacterial phyla identified in sample SRR33578315, with statistical classification (Very Low / Normal / Very High) relative to the healthy reference population.

Similarly, *Proteobacteria* (11.49%; *Very High*) was substantially increased, a finding often associated with inflammatory states and intestinal dysbiosis. No other phylum was detected in the sample: *Actinobacteria, Tenericutes, Verrucomicrobia, Cyanobacteria*, and *Fusobacteria* were all absent and classified as *Normal*, since for these low-prevalence phyla the reference distribution is zero-inflated and absence of the taxon corresponds to the reference point itself.

#### Associations with Clinical Conditions

The clinical association module identified bacterial alterations linked to health conditions across 11 clinical categories. The category *Gastrointestinal Diseases* accounted for the largest number of associations, with 22 alterations detected (13 positive and 9 negative). Other categories with a high number of associations included *Metabolic and Endocrine Diseases* (12), *Psychiatric and Neuropsychiatric Disorders* (10), and *Neoplasms and Neoplastic Syndromes* (9).

Of particular relevance, the *Metabolic and Endocrine Diseases* category—which encompasses type 2 Diabetes Mellitus—was among those with the highest number of associations, consistent with the diagnosis of the patient from whom the analyzed sample was derived.

Fig. 4 illustrates the output format of the association mod-ule for the condition *Type 2 Diabetes Mellitus*, demonstrating how the pipeline reports the direction of the association, the statistical positioning of the taxon, and the quantitative summary of the detected associations.

**Fig 4.**
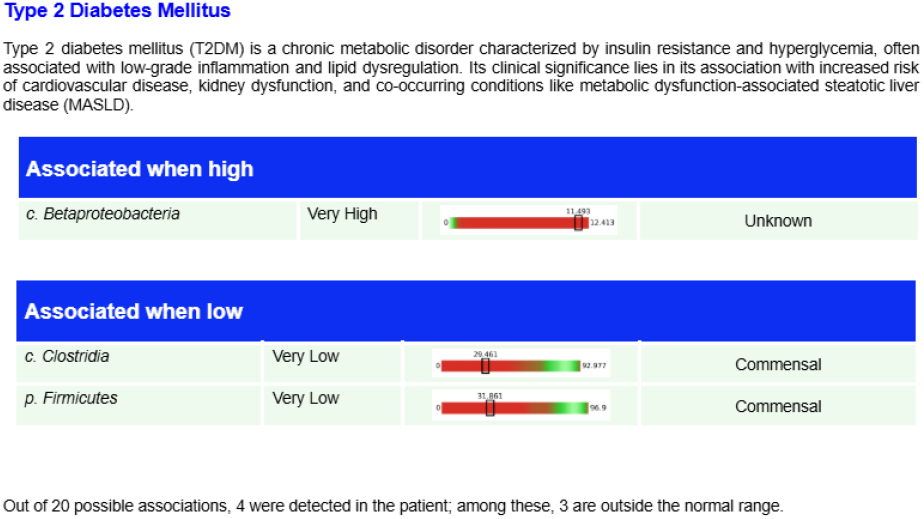
Example output of the clinical association module for Type 2 Diabetes Mellitus, showing taxa positively and negatively associated with the condition, their statistical classification, and a quantitative summary of detected associations.

### Generation of the Final Report

At the end of processing, the pipeline automatically generated a clinical report in .docx format, consolidating all results described in the previous sections. The report included diversity metrics, phylum-level abundances, enterotype classification, bacterial summaries by category, associations with clinical conditions, intervention recommendations, and a textual conclusion produced by the GPT-4o-mini model (14).

The conclusion section of the report generated for sample SRR33578315 illustrates the interpretive synthesis automatically produced by the pipeline, emphasizing dietary and lifestyle modifications—such as reducing high-sugar and high-fat processed foods and increasing fiber and fermented food intake—as strategies to support a more balanced gut microbiome. The entire execution was fully automated, with no manual intervention between steps. A complete example of the clinical report generated by the GTX-GUT pipeline for sample SRR33578315—including all sections, figures, and recommendations discussed in this section—is provided as Supplementary Material (Report S1).

## Discussion

The functional validation of the GTX-GUT pipeline using the ZymoBIOMICS mock community demonstrated its ability to accurately reproduce expected taxonomic profiles, confirming the robustness of the preprocessing and classification steps. The marked reduction in absolute error after applying BBDuk filtering highlights the importance of rigorous data preprocessing in minimizing biases introduced by sequencing artifacts. This finding is consistent with previous reports emphasizing that quality control and filtering are critical determinants of reliable metagenomic analysis (16).

Application of the pipeline to a human sample from a patient with type 2 Diabetes Mellitus provided further insights into its clinical relevance. The Shannon diversity index indicated medium-to-low alpha diversity, a pattern frequently associated with metabolic disorders and consistent with the liter-ature linking reduced microbial diversity to type 2 diabetes. Similarly, the Firmicutes/Bacteroidetes ratio, although technically within the clinically balanced range, was markedly reduced compared to healthy reference populations, reflecting a dominance of *Bacteroidetes*. This imbalance, together with the elevated abundance of *Proteobacteria*, suggests a dysbiotic profile characterized by inflammatory potential and metabolic disruption, in line with previous studies on diabetic cohorts (18).

The phylum-level analysis reinforced these observations, with *Firmicutes* showing a dramatic reduction relative to reference values and *Bacteroidetes* and *Proteobacteria* exhibiting substantial increases. Such shifts in microbial composition are well-documented hallmarks of gut dysbiosis and have been implicated in the pathophysiology of metabolic and inflammatory diseases. The consistency of these findings with established clinical associations underscores the pipeline’s capacity to capture biologically meaningful alterations in microbial communities.

The clinical association module further contextualized these taxonomic shifts by mapping them to health conditions across 11 categories. The predominance of associations with gastrointestinal, metabolic, and endocrine diseases is particularly relevant, as it aligns with the patient’s diagnosis of type 2 Diabetes Mellitus. This concordance between taxonomic alterations and clinical categories illustrates the potential of the pipeline to support translational applications, bridging microbiome research with clinical practice.

Finally, the automated generation of a comprehensive clinical report consolidating diversity metrics, taxonomic profiles, clinical associations, and interpretive conclusions represents a significant advance in usability and scalability. By eliminating the need for manual intervention, the pipeline ensures reproducibility and facilitates integration into clinical workflows. The inclusion of automated textual synthesis further enhances accessibility, enabling clinicians and researchers to rapidly interpret complex metagenomic data.

Taken together, these results demonstrate that GTX-GUT provides a reliable, reproducible, and clinically relevant framework for gut microbiome analysis. Its ability to validate against reference standards, detect disease-associated microbial signatures, and generate automated reports positions it as a valuable tool for both research and clinical applications (17). Future work should focus on expanding validation across diverse cohorts, integrating longitudinal data, and refining clinical association modules to strengthen predictive power and translational impact.

## Supporting information

Supplementary Report S1 - Example Clinical Report (SRR33578315)

## Bibliography

1. M. E. Walker, J. B. Simpson, and M. R. Redinbo. A structural metagenomics pipeline for examining the gut microbiome. Current Opinion in Structural Biology, 75:102416, 2022. doi: 10.1016/j.sbi.2022.102416.

2. J. Yepes-García and L. Falquet. 2Pipe starts with a question: Matching you with the correct pipeline for MAG reconstruction. mSystems, 11(2):e00844–25, 2026. doi: 10.1128/msystems.00844-25.

3. H. G. Lee, J. Y. Song, J. Yoon, Y. Chung, S.-K. Kwon, J. F. Kim, et al. A tool for metagenomic big data with fast and unified functional searches. Gut Microbes, 18(1):2611544, 2026. doi: 10.1080/19490976.2025.2611544.

4. F. Mölder, K. P. Jablonski, B. Letcher, M. B. Hall, C. H. Tomkins-Tinch, V. Sochat, et al. Sustainable data analysis with Snakemake. F1000Research, 10:33, 2021. doi: 10.12688/f1000research.29032.2.

5. G. M. Kurtzer, V. Sochat, and M. W. Bauer. Singularity: Scientific containers for mobility of compute. PLOS ONE, 12(5):e0177459, 2017. doi: 10.1371/journal.pone.0177459.

6. B. Bushnell. BBMap/BBDuk: A fast, versatile sequence trimming and filtering tool. Joint Genome Institute, 2014.

7. E. Bolyen, J. R. Rideout, M. R. Dillon, N. A. Bokulich, C. C. Abnet, G. A. Al-Ghalith, et al. Reproducible, interactive, scalable and extensible microbiome data science using QIIME 2. Nature Biotechnology, 37(8):852–857, 2019. doi: 10.1038/s41587-019-0209-9.

8. B. J. Callahan, P. J. McMurdie, M. J. Rosen, A. W. Han, A. J. A. Johnson, and S. P. Holmes. DADA2: High-resolution sample inference from Illumina amplicon data. Nature Methods, 13 (7):581–583, 2016. doi: 10.1038/nmeth.3869.

9. D. McDonald, M. N. Price, J. Goodrich, E. P. Nawrocki, T. Z. DeSantis, A. Probst, G. L. Andersen, R. Knight, and P. Hugenholtz. An improved Greengenes taxonomy with explicit ranks for ecological and evolutionary analyses of bacteria and archaea. The ISME Journal, 6(3):610–618, 2012. doi: 10.1038/ismej.2011.139.

10. C. E. Shannon. A mathematical theory of communication. The Bell System Technical Journal, 27(3):379–423, 1948.

11. J. A. Hartigan and P. M. Hartigan. The dip test of unimodality. The Annals of Statistics, 13 (1):70–84, 1985. doi: 10.1214/aos/1176346577.

12. M. Arumugam, J. Raes, E. Pelletier, D. Le Paslier, T. Yamada, D. R. Mende, et al. Enterotypes of the human gut microbiome. Nature, 473(7346):174–180, 2011. doi: 10.1038/nature09944.

13. L. Cheng, C. Qi, H. Zhuang, T. Fu, and X. Zhang. gutMDisorder: A comprehensive database for dysbiosis of the gut microbiota in disorders and interventions. Nucleic Acids Research, 48(D1):D554–D560, 2020. doi: 10.1093/nar/gkz843.

14. OpenAI. GPT-4o System Card. OpenAI, 2024.

15. Zymo Research. ZymoBIOMICS Microbial Community Standard (Catalog No. D6300): Certificate of Analysis. Zymo Research Corporation, 2024.

16. S. Mangul, L. S. Martin, B. L. Hill, A. K.-M. Lam, M. G. Distler, A. Zelikovsky, E. Eskin, and J. Flint. Systematic benchmarking of omics computational tools. Nature Communications, 10:1393, 2019. doi: 10.1038/s41467-019-09406-4.

17. M. A. Pita-Galeana, M. Ruhle, L. López-Vázquez, G. de Anda-Jáuregui, and E. Hernández-Lemus. Computational metagenomics: State of the art. International Journal of Molecular Sciences, 26(18):9206, 2025. doi: 10.3390/ijms26189206.

18. N. Larsen, F. K. Vogensen, F. W. J. van den Berg, D. S. Nielsen, A. S. Andreasen, B. K. Pedersen, et al. Gut microbiota in human adults with type 2 diabetes differs from non-diabetic adults. PLoS ONE, 5(2):e9085, 2010. doi: 10.1371/journal.pone.0009085.

19. R. E. Ley, P. J. Turnbaugh, S. Klein, and J. I. Gordon. Microbial ecology: Human gut microbes associated with obesity. Nature, 444(7122):1022–1023, 2006. doi: 10.1038/4441022a.

