## Supplementary Report S1 - Example Clinical Report (SRR33578315) for "GTX-GUT: A Standardized Metagenomic Workflow for Gut Microbiome Profiling and Clinical Associations"

+55 84 3113 5085

www.dnagtxbioinfo.com

### Gut Microbiota Metagenomics Report

| PATIENT | SAMPLE | REQUESTER |
|---|---|---|
| **Name:**<br>**ID:** SRR33578315<br>**Birthday:**<br>**Sex:** | Type:<br>Collection: day/month/year<br>Arrival: day/month/year | Requester:<br>Institution/company: |

#### 1. Introduction

This report presents a comprehensive set of recommendations aimed at optimizing your gut microbiome by addressing the abundance of various microorganisms. To promote a healthier gut environment, it is advised to reduce the intake of high-sugar and high-fat processed foods, which have been linked to the overgrowth of certain harmful bacteria such as Burkholderiales, Bacteroides, and Sutterella. Instead, increasing dietary fiber from fruits, vegetables, and whole grains is crucial, as it supports the growth of beneficial bacteria like Clostridia and Blautia. Incorporating fermented foods into your diet can further enhance microbial diversity and promote the growth of favorable species, including Parabacteroides distasonis and Oscillospira. However, it is important to note that some recommendations may seem contradictory, such as the need to limit high-fiber foods for certain bacteria like Lachnospira and Prevotella while simultaneously increasing fiber for others. Therefore, a balanced approach that focuses on moderation and variety in your diet, along with regular physical activity and the judicious use of antibiotics, will be essential in achieving a harmonious gut microbiome and overall well-being.t:

+55 84 3113 5085

www.dnagtxbioinfo.com

#### 2. Microbiota Diversity

Diversity is measured with two metrics: Firmicutes/Bacteroides Ratio (F/B Ratio) and Shannon Diversity Index (Shannon Index).

Patient F/B Ratio: **0.562**

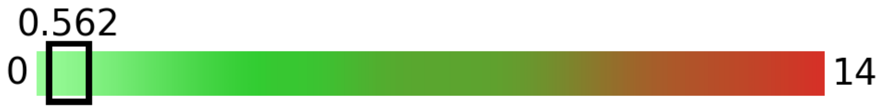


F/B < 1.5: Balanced

1.5 ≤ F/B ≤ 3: Imbalanced microbiota

F/B > 3: Unfavorable microbiota

The F/B Ratio is a critical metric that measures the balance between the two dominant groups of bacteria in your gut: Firmicutes and Bacteroidetes. A balanced ratio (below 1.5) indicates a healthy, diverse microbiome typically associated with high-fiber diets and efficient metabolic function. When this ratio climbs between 1.5 and 3.0, it signals an imbalance often caused by a "Western diet" high in processed sugars and fats. An unfavorable ratio (above 3.0) is frequently linked to obesity and metabolic issues, as Firmicutes are highly efficient at extracting and storing extra calories from your food. By tracking this number, you can see a direct "snapshot" of how your dietary choices are influencing your gut's ability to manage energy and inflammation. Improving your score is usually as simple as increasing plant diversity and fiber to help your "lean-profile" Bacteroidetes thrive again.

Patient Shannon Index: **2.571**

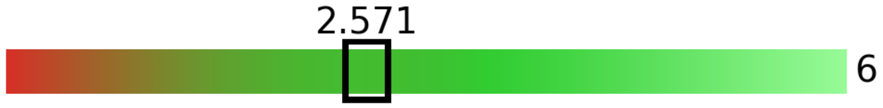


Shannon < 3: Limited diversity

3 ≤ Shannon ≤ 4: Medium diversity

Shannon > 4: Good diversity

The Shannon Index is the primary measure of your gut's biodiversity, calculating both the number of different bacterial species present and how evenly they are distributed. A High Diversity score (above 4.0) indicates a resilient ecosystem that is better equipped to fight off infections and regulate your immune system. If your score falls in the Medium range (3.0 to 4.0), your gut is functional but could benefit from a wider variety of plant-based foods to increase its stability. A Limited score (below 3.0) is a warning sign of a "thinned out" microbiome, often linked to high stress, poor diet, or recent antibiotic treatments. Think of your gut like a rainforest: the more diverse the inhabitants, the healthier and more productive the entire environment becomes. By tracking this index over time, you can see the direct impact that adding diverse fibers and fermented foods has on your internal ecosystem.


BRAZIL | UAE | USA
Av. Senador Salgado Filho, 3.000 - Room A402 – CEP: 59.078-970 – Natal / RN
Av. Paulista, 1.636 - Conjunto 301 – CEP: 13.102-000 – São Paulo / SP
DNA GTx

### 3. Bacterial Summary

Bacteria in this report can be classified into 3 categories:

**Pathogenic Bacteria:** Harmful species which cause infections and contagious diseases;

**Commensal Bacteria:** Species that naturally live in the human gut without causing harm and providing benefits to the host;

**Bacteria With Uncertain Effects:** Bacteria which need further research to improve scientific understanding of their roles in the human microbiota;

##### Bacteria by Abundance Levels

Abundance levels are compared to populational data and classified as very low, normal or very high. Abundances are very low or very high when they are statistical outliers (very statistically uncommon).

Diseases are described in detail only when they are associated with bacteria showing abnormal abundance. The related bacteria are displayed exclusively as evidence of the association.

| Classification | Abundance Levels | | |
|---|---|---|---|
| | Very Low | Normal | Very High |
| Pathogenic Bacteria | 0 | 0 | 1 |
| Commensal Bacteria | 2 | 15 | 12 |
| Bacteria With Uncertain Effects | 2 | 3 | 6 |

##### Enterotype: Enterotype 1 (Bacteroides-dominant)

+55 84 3113 5085

www.dnagtxbioinfo.com

#### Abundance of Bacterial Phylum

These are the main types of bacteria, which include most of the species of bacteria found in the human gut.

| | | | |
|---|---|---|---|
| *Firmicutes* | 0 – 96.90 | Very Low | Help digest food, extract energy, and maintain gut health |
| *Bacteroidetes* | 0 – 61.177 | Very High | Break down complex carbs, produce beneficial fats, and support immunity |
| *Actinobacteria* | 0 – 4.946 | Normal | Aid digestion, protect against harmful bacteria, and boost gut health |
| *Proteobacteria* | 0 – 12.413 | Very High | Can cause infections but also help with nutrient processing in small amounts |
| *Tenericutes* | 0 – 0.720 | Normal | Some are harmless, while others may cause diseases in the gut or elsewhere |
| *Verrucomicrobia* | 0 – 0.480 | Normal | Strengthen the gut lining and help regulate mucus production |
| *Cyanobacteria* | 0 – 0.250 | Normal | Rare in the gut; some can produce toxins, others may have minor roles |
| *Fusobacteria* | 0 – 0.250 | Normal | Often linked to infections but can also be part of a balanced microbiome |

+55 84 3113 5085

www.dnagtxbioinfo.com

### 4. Disease Associations

#### Bacterial associations grouped by clinical category

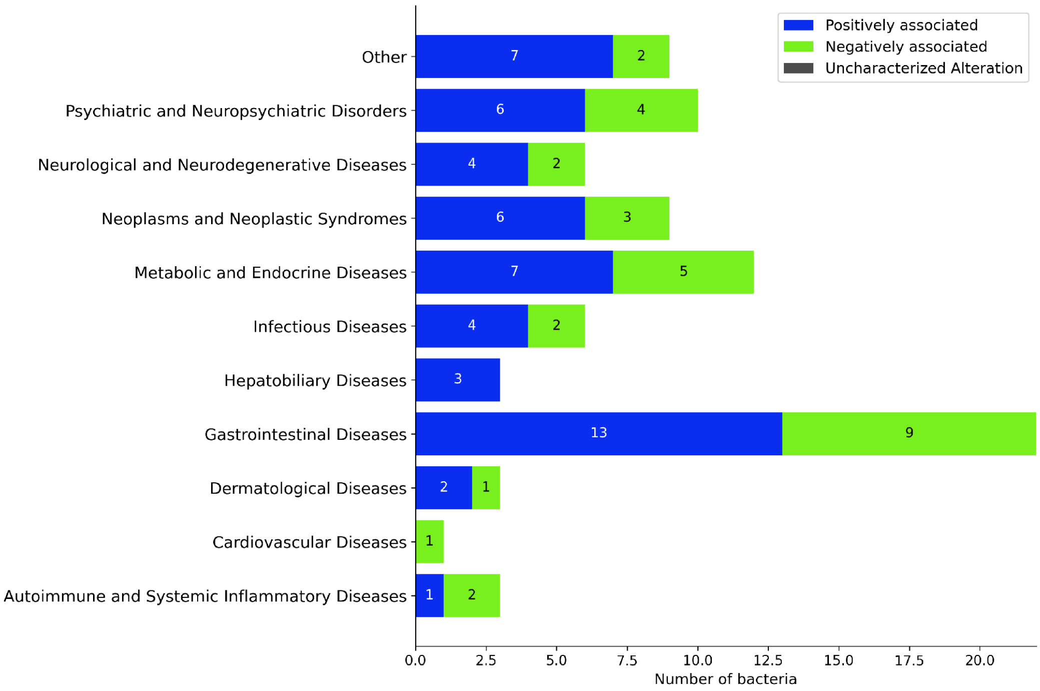


#### Altered bacteria in each clinical category

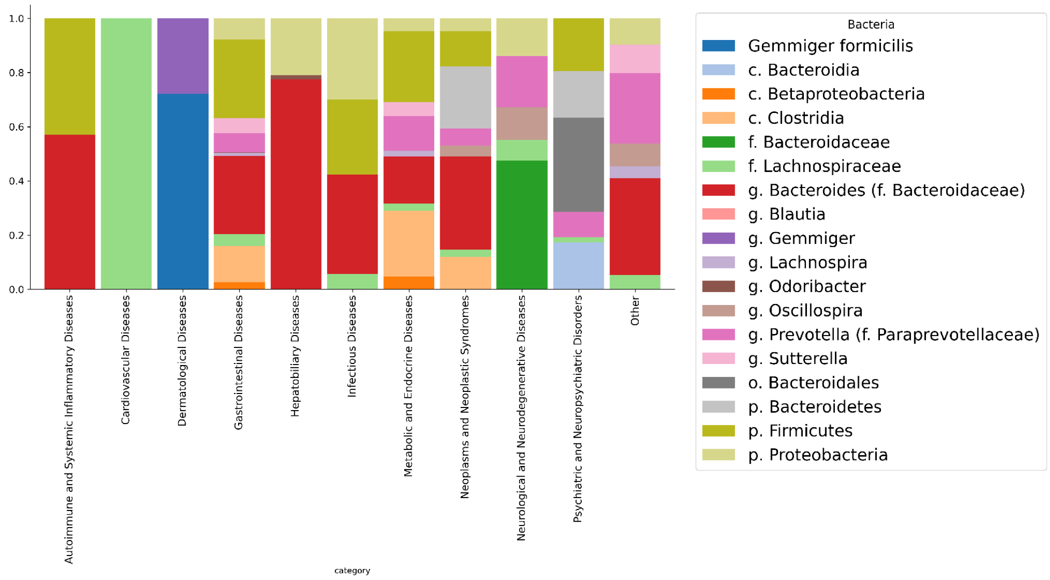

### Autoimmune and Systemic Inflammatory Diseases

#### Acute Anterior Uveitis

Acute anterior uveitis is an inflammatory condition affecting the front part of the uvea, which can lead to significant visual impairment; accurate identification of its viral causes is crucial for effective treatment.

| Associated when low | | | |
|---|---|---|---|
| *g. Blautia* | Very Low | 0 — 0 — 30.776 | Commensal |

Out of 7 possible associations, 2 were detected in the patient; among these, 1 is outside the normal range.

#### Systemic Lupus Erythematosus

Systemic lupus erythematosus (SLE) is a chronic autoimmune disorder marked by immune system dysregulation, resulting in widespread inflammation and tissue damage. Its pathogenesis involves the dysfunction of tolerogenic dendritic cells and regulatory T cells, highlighting the importance of immune balance in disease progression.

| Associated when low | | | |
|---|---|---|---|
| *p. Firmicutes* | Very Low | 0 — 31.861 — 96.9 | Commensal |

Out of 1 possible associations, 1 was detected in the patient; among these, 1 is outside the normal range.

#### Type 1 Diabetes Mellitus

Type 1 diabetes mellitus is an autoimmune condition characterized by the destruction of insulin-producing pancreatic beta cells, leading to absolute insulin deficiency. Its clinical significance lies in the increased risk of hypoglycemia and the need for exogenous insulin to maintain glycemic control.

| Associated when high | | | |
|---|---|---|---|
| *g. Bacteroides (f. Bacteroidaceae)* | Very High | 0 — 42.3 — 45.684 | Commensal |

Out of 31 possible associations, 6 were detected in the patient; among these, 1 is outside the normal range.

### Cardiovascular Diseases

#### Unstable Angina

Unstable angina is a clinical syndrome characterized by unpredictable chest pain due to transient myocardial ischemia, often indicating significant coronary artery disease and an increased risk of myocardial infarction. Its presence necessitates urgent medical evaluation and intervention to prevent severe cardiovascular events.

| Associated when low | | | |
|---|---|---|---|
| *f. Lachnospiraceae* | Very Low | 0 · 6.439 · 67.829 | Unknown |

Out of 6 possible associations, 1 was detected in the patient; among these, 1 is outside the normal range.

#### Dermatological Diseases

##### Acne Vulgaris

Acne vulgaris is a common skin condition characterized by the presence of comedones, papules, and pustules due to the obstruction of hair follicles and inflammation, often linked to hormonal changes and bacterial proliferation. Clinically, it significantly impacts patients' quality of life and may require comprehensive management strategies, including topical treatments and adjunctive therapies.

| Associated when low | | | |
|---|---|---|---|
| *g. Blautia* | Very Low | 0 · 0 · 30.776 | Commensal |

Out of 18 possible associations, 4 were detected in the patient; among these, 1 is outside the normal range.

##### Eczema

Eczema, or atopic dermatitis, is a chronic inflammatory skin condition characterized by pruritic, erythematous lesions and is associated with immune dysregulation. Its clinical significance lies in its impact on quality of life and its comorbidity with mental health disorders, such as depression.

| Associated when high | | | |
|---|---|---|---|
| *g. Gemmiger* | Very High | 0 · 7.558 · 8.162 | Commensal |
| *Gemmiger formicilis* | Very High | 0 · 19.538 · 21.101 | Commensal |

Out of 66 possible associations, 9 were detected in the patient; among these, 2 are outside the normal range.

#### Gastrointestinal Diseases

+55 84 3113 5085

www.dnagtxbioinfo.com

#### Celiac Disease

Celiac disease is an autoimmune disorder characterized by an inappropriate immune response to gluten, leading to intestinal damage and malabsorption. Its clinical significance lies in the potential for severe complications if left untreated, including nutritional deficiencies and increased risk of other autoimmune conditions.

| Associated when low | | | |
|---|---|---|---|
| *p. Firmicutes* | Very Low | 0 — 31.861 — 96.9 | Commensal |

Out of 6 possible associations, 1 was detected in the patient; among these, 1 is outside the normal range.

#### Crohn Disease

Crohn's disease (CD) is a chronic inflammatory bowel disorder characterized by inflammation and potential fibrosis of the intestinal tract, leading to complications such as strictures. The study highlights dipeptidyl peptidase 4 (DPP4) as a significant driver of intestinal fibrosis in CD, suggesting it as a novel therapeutic target for managing this condition.

| Associated when high | | | |
|---|---|---|---|
| *g. Bacteroides (f. Bacteroidaceae)* | Very High | 0 — 42.3 — 45.684 | Commensal |
| *g. Odoribacter* | Very High | 0 — 0.787 — 0.85 | Commensal |
| *g. Prevotella (f. Paraprevotellaceae)* | Very High | 0 — 15.471 — 16.709 | Unknown |
| *g. Sutterella* | Very High | 0 — 12.441 — 13.437 | Unknown |
| *p. Proteobacteria* | Very High | 0 — 11.493 — 12.413 | Commensal |

| Associated when low | | | |
|---|---|---|---|
| *p. Firmicutes* | Very Low | 0 — 31.861 — 96.9 | Commensal |

Out of 90 possible associations, 15 were detected in the patient; among these, 6 are outside the normal range.

#### Diarrhea

Diarrhea is characterized by increased frequency and fluidity of bowel movements, often resulting from infections, which can lead to dehydration and electrolyte imbalances. Its clinical significance lies in the potential for severe complications, particularly in vulnerable populations, necessitating effective treatment strategies.



BRAZIL | UAE | USA
Av. Senador Salgado Filho, 3.000 - Room A402 – CEP: 59.078-970 – Natal / RN
Av. Paulista, 1.636 - Conjunto 301 – CEP: 13.102-000 – São Paulo / SP
DNA GTx

+55 84 3113 5085

www.dnagtxbioinfo.com

| Associated when low | | | |
|---|---|---|---|
| *f. Lachnospiraceae* | Very Low | 0 6.439 67.829 | Unknown |

Out of 17 possible associations, 8 were detected in the patient; among these, 1 is outside the normal range.

#### Dysentery

Dysentery is an inflammatory disorder of the intestines, primarily characterized by severe diarrhea with blood and mucus, often caused by infectious agents such as bacteria or parasites. Clinically significant, it can lead to dehydration and systemic complications, particularly in vulnerable populations.

| Associated when high | | | |
|---|---|---|---|
| *g. Sutterella* | Very High | 0 12.441 13.437 | Unknown |

Out of 16 possible associations, 3 were detected in the patient; among these, 1 is outside the normal range.

#### First-Degree Relatives Of Children With Crohn’s Disease

First-degree relatives of children with Crohn's disease are individuals, such as parents or siblings, who share a genetic predisposition to inflammatory bowel disease (IBD), potentially increasing their risk of developing the condition. Understanding their perceptions regarding predictive testing and prevention is crucial for improving early intervention strategies and managing familial IBD risk.

| Associated when low | | | |
|---|---|---|---|
| *f. Lachnospiraceae* | Very Low | 0 6.439 67.829 | Unknown |

Out of 12 possible associations, 2 were detected in the patient; among these, 1 is outside the normal range.

#### Irritable Bowel Syndrome

Irritable bowel syndrome (IBS) is a functional gastrointestinal disorder characterized by abdominal pain and altered bowel habits, often accompanied by bloating. Its clinical significance lies in its impact on quality of life and the challenges it poses for diagnosis and management, as it lacks identifiable organic causes.

| Associated when high | | | |
|---|---|---|---|
| *g. Lachnospira* | Very High | 0 5.328 5.754 | Commensal |

+55 84 3113 5085

www.dnagtxbioinfo.com

Out of 36 possible associations, 3 were detected in the patient; among these, 1 is outside the normal range.

#### Necrotizing Enterocolitis

Necrotizing enterocolitis (NEC) is a severe gastrointestinal condition primarily affecting premature infants, characterized by inflammation and necrosis of the intestinal tissue, leading to significant morbidity and mortality. Its clinical significance lies in its association with preterm birth and the high risk of complications, necessitating prompt diagnosis and management.

| Associated when high | | | |
|---|---|---|---|
| *g. Bacteroides (f. Bacteroidaceae)* | Very High | 0 – 42.3 – 45.684 | Commensal |

| Associated when low | | | |
|---|---|---|---|
| *c. Clostridia* | Very Low | 0 – 29.461 – 92.977 | Commensal |

Out of 16 possible associations, 5 were detected in the patient; among these, 4 are outside the normal range.

#### Necrotizing Enterocolitis In Premature Infant

Colitis is an inflammatory condition of the colon characterized by symptoms such as abdominal pain and diarrhea, often resulting from dysregulated immune responses. Its clinical significance lies in the potential for severe complications and the need for effective therapeutic interventions to restore intestinal homeostasis.

| Associated when high | | | |
|---|---|---|---|
| *p. Proteobacteria* | Very High | 0 – 11.493 – 12.413 | Commensal |

| Associated when low | | | |
|---|---|---|---|
| *p. Firmicutes* | Very Low | 0 – 31.861 – 96.9 | Commensal |

#### Severe Acute Malnutrition And Diarrhea

Diarrhea is characterized by increased frequency and fluidity of bowel movements, often resulting from infections, which can lead to dehydration and electrolyte imbalances. Its clinical significance lies in the potential for severe complications, particularly in vulnerable populations, necessitating effective treatment strategies.


BRAZIL | UAE | USA
Av. Senador Salgado Filho, 3.000 - Room A402 – CEP: 59.078-970 – Natal / RN
Av. Paulista, 1.636 - Conjunto 301 – CEP: 13.102-000 – São Paulo / SP
DNA GTx

+55 84 3113 5085

www.dnagtxbioinfo.com

| Associated when low | | | |
|---|---|---|---|
| *f. Lachnospiraceae* | Very Low | 6.439 (0 – 67.829) | Unknown |

##### Ulcerative Colitis

Ulcerative colitis (UC) is a chronic inflammatory bowel disease characterized by inflammation and ulceration of the colonic mucosa, leading to symptoms such as diarrhea and abdominal pain. Its clinical significance lies in its association with increased risk of colorectal cancer and the potential for systemic complications.

| Associated when high | | | |
|---|---|---|---|
| *g. Bacteroides (f. Bacteroidaceae)* | Very High | 42.3 (0 – 45.684) | Commensal |
| *g. Prevotella (f. Paraprevotellaceae)* | Very High | 15.471 (0 – 16.709) | Unknown |
| *c. Betaproteobacteria* | Very High | 11.493 (0 – 12.413) | Unknown |
| *p. Proteobacteria* | Very High | 11.493 (0 – 12.413) | Commensal |

| Associated when low | | | |
|---|---|---|---|
| *c. Clostridia* | Very Low | 29.461 (0 – 92.977) | Commensal |
| *p. Firmicutes* | Very Low | 31.861 (0 – 96.9) | Commensal |

Out of 63 possible associations, 8 were detected in the patient; among these, 6 are outside the normal range.

#### Hepatobiliary Diseases

##### Cholesterol Gallstones

Cholesterol gallstones are solid particles that form in the gallbladder due to supersaturation of cholesterol in bile, often associated with metabolic disorders like obesity and insulin resistance. Their clinical significance lies in their potential to cause biliary colic, cholecystitis, and other complications, highlighting the importance of understanding their pathogenesis.

| Associated when high | | | |
|---|---|---|---|
| *p. Proteobacteria* | Very High | 0 – 11.493 – 12.413 | Commensal |

Out of 4 possible associations, 2 were detected in the patient; among these, 1 is outside the normal range.

##### Hbv-Related Hepatocellular Carcinoma

HBV-related hepatocellular carcinoma (HCC) is a liver cancer that arises from chronic infection with the hepatitis B virus, contributing significantly to global cancer morbidity and mortality. Its genomic landscape shows similarities to other HCC etiologies, but specific mutations may influence treatment strategies and patient outcomes.

| Associated when high | | | |
|---|---|---|---|
| *g. Bacteroides (f. Bacteroidaceae)* | Very High | 0 – 42.3 – 45.684 | Commensal |

Out of 2 possible associations, 1 was detected in the patient; among these, 1 is outside the normal range.

##### Non-Alcoholic Fatty Liver Disease

Non-alcoholic fatty liver disease (NAFLD) is a condition characterized by excessive fat accumulation in the liver in individuals who consume little to no alcohol, often linked to obesity and metabolic syndrome. Clinically significant, NAFLD can progress to more severe liver diseases, including steatohepatitis, fibrosis, and cirrhosis, thereby impacting overall metabolic health.

| Associated when high | | | |
|---|---|---|---|
| *g. Odoribacter* | Very High | 0 – 0.787 – 0.85 | Commensal |

Out of 20 possible associations, 4 were detected in the patient; among these, 1 is outside the normal range.

#### Infectious Diseases

##### Caliciviridae Infections

Caliciviridae infections, primarily caused by noroviruses, are significant contributors to acute gastroenteritis globally, particularly in children. The recent emergence and predominance of specific genotypes, such as GII.17[P17], highlight the evolving genetic diversity and public health impact of these viral infections.

| Associated when high | | | |
|---|---|---|---|
| *p. Proteobacteria* | Very High | 0 – 11.493 – 12.413 | Commensal |

Out of 2 possible associations, 2 were detected in the patient; among these, 1 is outside the normal range.

#### Clostridium Infections

Clostridium infections, primarily caused by Clostridium difficile, are significant gastrointestinal diseases characterized by antibiotic-associated diarrhea and colitis, particularly in individuals with disrupted gut microbiota. These infections can lead to severe complications, including toxic megacolon and increased mortality in vulnerable populations.

| Associated when high | | | |
|---|---|---|---|
| *g. Bacteroides (f. Bacteroidaceae)* | Very High | 0 — 42.3 — 45.684 | Commensal |
| *p. Proteobacteria* | Very High | 0 — 11.493 — 12.413 | Commensal |

| Associated when low | | | |
|---|---|---|---|
| *f. Lachnospiraceae* | Very Low | 0 — 6.439 — 67.829 | Unknown |
| *p. Firmicutes* | Very Low | 0 — 31.861 — 96.9 | Commensal |

Out of 12 possible associations, 6 were detected in the patient; among these, 4 are outside the normal range.

#### Hiv

HIV infections are caused by the human immunodeficiency virus (HIV), which attacks the immune system, specifically CD4+ T cells, leading to progressive immune deficiency. This condition, if untreated, can result in acquired immunodeficiency syndrome (AIDS), characterized by severe opportunistic infections and malignancies due to the body's inability to mount an effective immune response. Clinically, HIV infections are significant as they pose a major global health challenge, affecting millions and leading to substantial morbidity and mortality. The virus is primarily transmitted through unprotected sexual contact, sharing of contaminated needles, and from mother to child during childbirth or breastfeeding. Effective antiretroviral therapy (ART) can manage HIV infections, allowing individuals to maintain a near-normal life expectancy and significantly reduce the risk of transmission. Public health strategies, including education, access to testing, and treatment, are crucial in controlling the spread of HIV and

| Associated when high | | | |
|---|---|---|---|
| *p. Proteobacteria* | Very High | 0 — 11.493 — 12.413 | Commensal |

Out of 10 possible associations, 7 were detected in the patient; among these, 1 is outside the normal range.

### Metabolic and Endocrine Diseases

#### Gestational Diabetes

Gestational diabetes mellitus (GDM) is a form of glucose intolerance that occurs during pregnancy, characterized by elevated blood sugar levels that can affect both maternal and fetal health. Its biological significance lies in its association with increased risks of complications such as preeclampsia, cesarean delivery, and long-term metabolic disorders for both mother and child.

| Associated when low | | | |
|---|---|---|---|
| *p. Firmicutes* | Very Low | 0 – 31.861 – 96.9 | Commensal |

Out of 2 possible associations, 2 were detected in the patient; among these, 1 is outside the normal range.

#### Hepatolenticular Degeneration

Hepatolenticular degeneration, also known as Wilson's disease, is a genetic disorder characterized by excessive copper accumulation in the liver and brain, leading to hepatic and neurological dysfunction. Its clinical significance lies in the potential for severe liver damage and neurological impairment if left untreated.

| Associated when high | | | |
|---|---|---|---|
| *g. Bacteroides (f. Bacteroidaceae)* | Very High | 0 – 42.3 – 45.684 | Commensal |
| *g. Lachnospira* | Very High | 0 – 5.328 – 5.754 | Commensal |

Out of 7 possible associations, 3 were detected in the patient; among these, 2 are outside the normal range.

#### Obesity

Obesity is a complex metabolic disorder characterized by excessive fat accumulation that poses significant health risks, including cardiovascular diseases, diabetes, and certain cancers. Its clinical significance lies in its association with increased morbidity and mortality, necessitating effective management strategies.

| Associated when high | | | |
|---|---|---|---|
| *g. Prevotella (f. Paraprevotellaceae)* | Very High | 0 – 15.471 – 16.709 | Unknown |
| *g. Sutterella* | Very High | 0 – 12.441 – 13.437 | Unknown |
| *p. Proteobacteria* | Very High | 0 – 11.493 – 12.413 | Commensal |

| Associated when low | | | |
|---|---|---|---|
| *c. Clostridia* | Very Low | 0 – 29.461 – 92.977 | Commensal |

Out of 42 possible associations, 11 were detected in the patient; among these, 4 are outside the normal range.

#### Phenylketonuria

Phenylketonuria (PKU) is a genetic disorder caused by a deficiency in the enzyme phenylalanine hydroxylase, leading to the accumulation of phenylalanine, which can cause intellectual disability and neurological issues if untreated. Early diagnosis and dietary management are crucial to prevent severe cognitive impairment and other complications.

| Associated when high | | | |
|---|---|---|---|
| *g. Prevotella (f. Paraprevotellaceae)* | Very High | 0 — 15.471 — 16.709 | Unknown |

| Associated when low | | | |
|---|---|---|---|
| *f. Lachnospiraceae* | Very Low | 0 — 6.439 — 67.829 | Unknown |

Out of 11 possible associations, 7 were detected in the patient; among these, 2 are outside the normal range.

#### Type 2 Diabetes Mellitus

Type 2 diabetes mellitus (T2DM) is a chronic metabolic disorder characterized by insulin resistance and hyperglycemia, often associated with low-grade inflammation and lipid dysregulation. Its clinical significance lies in its association with increased risk of cardiovascular disease, kidney dysfunction, and co-occurring conditions like metabolic dysfunction-associated steatotic liver disease (MASLD).

| Associated when high | | | |
|---|---|---|---|
| *c. Betaproteobacteria* | Very High | 0 — 11.493 — 12.413 | Unknown |

| Associated when low | | | |
|---|---|---|---|
| *c. Clostridia* | Very Low | 0 29.461 92.977 | Commensal |
| *p. Firmicutes* | Very Low | 0 31.861 96.9 | Commensal |

Out of 20 possible associations, 4 were detected in the patient; among these, 3 are outside the normal range.

### Neoplasms and Neoplastic Syndromes

#### Colorectal Neoplasms

Colorectal neoplasms are abnormal growths in the colon or rectum, which can be benign or malignant, with colorectal cancer being a leading cause of cancer-related morbidity and mortality. Their biological significance lies in the potential for metastasis and the need for early detection and intervention to improve patient outcomes.

| Associated when high | | | |
|---|---|---|---|
| *g. Bacteroides (f. Bacteroidaceae)* | Very High | 0 42.3 45.684 | Commensal |
| *g. Oscillospira* | Very High | 0 9.932 10.726 | Commensal |
| *g. Prevotella (f. Paraprevotellaceae)* | Very High | 0 15.471 16.709 | Unknown |

| Associated when low | | | |
|---|---|---|---|
| *c. Clostridia* | Very Low | 0 29.461 92.977 | Commensal |
| *p. Firmicutes* | Very Low | 0 31.861 96.9 | Commensal |

Out of 108 possible associations, 12 were detected in the patient; among these, 5 are outside the normal range.

#### Colorectal Adenomatous Polyposis, Autosomal Recessive

Colorectal Adenomatous Polyposis, Autosomal Recessive (CAPAR) is a genetic condition characterized by the development of multiple adenomatous polyps in the colon and rectum, leading to an increased risk of colorectal cancer. It is caused by biallelic mutations in specific genes, such as NTHL1 and MUTYH, which are involved in DNA repair mechanisms. The clinical significance of CAPAR lies in its association with the early onset of colorectal neoplasms, necessitating regular surveillance and potential prophylactic interventions. Recent studies have identified colibactin, a genotoxin produced by certain strains of Escherichia coli, as a contributing factor to mutational processes in colorectal tumors associated with CAPAR, highlighting the interplay between genetic predisposition and microbial influences in cancer development. Understanding these mechanisms is crucial for improving screening strategies and therapeutic approaches for affected

+55 84 3113 5085

www.dnagtxbioinfo.com

| Associated when high | | | |
|---|---|---|---|
| *p. Bacteroidetes* | Very High | 0 – 56.646 – 61.177 | Commensal |
| *p. Proteobacteria* | Very High | 0 – 11.493 – 12.413 | Commensal |

Out of 8 possible associations, 3 were detected in the patient; among these, 2 are outside the normal range.

#### Hereditary Mixed Polyposis Syndrome

Hereditary mixed polyposis syndrome is a genetic condition characterized by the presence of various types of polyps in the gastrointestinal tract, primarily associated with GREM1 and BMPR1A gene variants. Its clinical significance lies in an increased risk of colorectal cancer and the need for vigilant surveillance and management strategies.

| Associated when high | | | |
|---|---|---|---|
| *g. Bacteroides (f. Bacteroidaceae)* | Very High | 0 – 42.3 – 45.684 | Commensal |

| Associated when low | | | |
|---|---|---|---|
| *f. Lachnospiraceae* | Very Low | 0 – 6.439 – 67.829 | Unknown |

Out of 4 possible associations, 3 were detected in the patient; among these, 2 are outside the normal range.

### Neurological and Neurodegenerative Diseases

#### Drug Resistant Epilepsy

Drug-resistant epilepsy is a form of epilepsy that does not respond to conventional antiepileptic medications, significantly impacting patients' quality of life and often necessitating surgical intervention. It is clinically significant as it may be associated with underlying structural or genetic abnormalities, such as focal cortical dysplasia, which can be targeted for precision therapies.

| Associated when high | | | |
|---|---|---|---|
| *g. Prevotella (f. Paraprevotellaceae)* | Very High | 0 – 15.471 – 16.709 | Unknown |
| *p. Proteobacteria* | Very High | 0 – 11.493 – 12.413 | Commensal |

Out of 3 possible associations, 2 were detected in the patient; among these, 2 are outside the normal range.

+55 84 3113 5085

www.dnagtxbioinfo.com

#### Multiple Sclerosis

Multiple sclerosis (MS) is a chronic autoimmune disorder characterized by the demyelination of neurons in the central nervous system, leading to neurological dysfunction. Its pathogenesis involves complex interactions between genetic predisposition, environmental factors, and immune system dysregulation.

| Associated when low | | | |
|---|---|---|---|
| *f. Lachnospiraceae* | Very Low | 6.439 (0 – 67.829) | Unknown |

Out of 15 possible associations, 3 were detected in the patient; among these, 1 is outside the normal range.

#### Parkinson Disease

Parkinson's disease (PD) is a progressive neurodegenerative disorder characterized by the loss of dopaminergic neurons in the substantia nigra, leading to motor and non-motor symptoms. Its clinical significance lies in the impact on movement control and quality of life, as well as its association with altered hemodynamics and metabolism in the brain.

| Associated when high | | | |
|---|---|---|---|
| *g. Oscillospira* | Very High | 9.932 (0 – 10.726) | Commensal |

| Associated when low | | | |
|---|---|---|---|
| *g. Blautia* | Very Low | 0 (0 – 30.776) | Commensal |

Out of 41 possible associations, 7 were detected in the patient; among these, 2 are outside the normal range.

#### Rett Syndrome

Rett syndrome is a rare X-linked neurodevelopmental disorder caused by mutations in the MECP2 gene, leading to severe impairments in communication, motor skills, and autonomic functions. Clinically, it is characterized by progressive loss of purposeful hand skills, gait abnormalities, and dysphagia, significantly impacting quality of life.

| Associated when high | | | |
|---|---|---|---|
| *f. Bacteroidaceae* | Very High | 39.077 (0 – 42.204) | Commensal |

Out of 2 possible associations, 1 was detected in the patient; among these, 1 is outside the normal range.


BRAZIL | UAE | USA
Av. Senador Salgado Filho, 3.000 - Room A402 – CEP: 59.078-970 – Natal / RN
Av. Paulista, 1.636 - Conjunto 301 – CEP: 13.102-000 – São Paulo / SP
DNA GTx

### Psychiatric and Neuropsychiatric Disorders

#### Anorexia Nervosa

Anorexia nervosa (AN) is a severe psychiatric disorder marked by extreme caloric restriction and significant weight loss, leading to detrimental physiological changes, particularly in skeletal muscle. Its clinical significance lies in the persistent alterations in muscle proteomics and function, even after weight restoration, highlighting the complexity of recovery and the need for targeted interventions.

| Associated when high | | | |
|---|---|---|---|
| *c. Bacteroidia* | Very High | 0 56.646 61.177 | Commensal |
| *o. Bacteroidales* | Very High | 0 56.646 61.177 | Commensal |
| *p. Bacteroidetes* | Very High | 0 56.646 61.177 | Commensal |

| Associated when low | | | |
|---|---|---|---|
| *p. Firmicutes* | Very Low | 0 31.861 96.9 | Commensal |

Out of 24 possible associations, 11 were detected in the patient; among these, 5 are outside the normal range.

#### Attention Deficit Disorder With Hyperactivity

Attention deficit hyperactivity disorder (ADHD) is a neurodevelopmental disorder characterized by persistent patterns of inattention, hyperactivity, and impulsivity, which can significantly impair functioning and development. Its biological significance lies in the dysregulation of neuroactive steroids and other neurodevelopmental pathways, particularly influenced by maternal factors such as obesity, which may increase the risk of ADHD in offspring.

| Associated when high | | | |
|---|---|---|---|
| *g. Prevotella (f. Paraprevotellaceae)* | Very High | 0 15.471 16.709 | Unknown |

| Associated when low | | | |
|---|---|---|---|
| *p. Firmicutes* | Very Low | 0 31.861 96.9 | Commensal |

Out of 6 possible associations, 3 were detected in the patient; among these, 2 are outside the normal range.

#### Depression

Depression is a common mental health disorder characterized by persistent sadness, loss of interest, and functional impairment, often linked to neurobiological changes and psychosocial stressors. Its clinical significance lies in its association with increased morbidity, impaired quality of life, and heightened risk of suicide.

| Associated when high | | | |
|---|---|---|---|
| *o. Bacteroidales* | Very High | 0 — 56.646 — 61.177 | Commensal |
| *g. Prevotella (f. Paraprevotellaceae)* | Very High | 0 — 15.471 — 16.709 | Unknown |

| Associated when low | | | |
|---|---|---|---|
| *g. Blautia* | Very Low | 0 — 0 — 30.776 | Commensal |
| *f. Lachnospiraceae* | Very Low | 0 — 6.439 — 67.829 | Unknown |

Out of 25 possible associations, 6 were detected in the patient; among these, 4 are outside the normal range.

### Other

#### Anti-Bacterial Agents(Vancomycin And Metronidazole)

Vancomycin and metronidazole are broad-spectrum antibiotics used to treat infections caused by Clostridioides difficile, particularly in cases of severe or recurrent disease. Their clinical significance lies in their ability to manage CDI symptoms and prevent complications, especially in immunocompromised patients.

| Associated when high | | | |
|---|---|---|---|
| *g. Bacteroides (f. Bacteroidaceae)* | Very High | 0 — 42.3 — 45.684 | Commensal |
| *g. Oscillospira* | Very High | 0 — 9.932 — 10.726 | Commensal |

Out of 11 possible associations, 2 were detected in the patient; among these, 2 are outside the normal range.

#### Antiretroviral Therapy,highly Active; Hiv Infections

Highly Active Antiretroviral Therapy (HAART) is a comprehensive treatment regimen for individuals living with HIV/AIDS, employing a combination of at least three antiretroviral drugs from different classes to suppress viral replication and improve immune function. The clinical significance of HAART lies in its ability to transform HIV from a fatal disease into a manageable chronic condition, significantly reducing AIDS-related morbidity and mortality.

In the context of Ethiopian health facilities, the expansion of HAART has led to a notable decrease in AIDS-related deaths; however, the survival rates among patients on therapy exhibit considerable variability. Factors such as socio-demographic characteristics, clinical stage of HIV, and adherence to treatment are critical determinants of early mortality among patients. For instance, advanced clinical stages (WHO stages III/IV) and poor adherence to therapy have been identified as significant predictors of increased mortality risk.

This underscores the importance of early enrollment in

| Associated when high | | | |
|---|---|---|---|
| *g. Prevotella (f. Paraprevotellaceae)* | Very High | 0 15.471 16.709 | Unknown |

#### Blastocystis Infections

Blastocystis infections are caused by the protozoan parasite Blastocystis spp., which is the most commonly identified protozoan in humans. The clinical significance of these infections is debated, as many individuals remain asymptomatic. However, Blastocystis has been linked to gastrointestinal symptoms such as diarrhea and abdominal pain, as well as extraintestinal manifestations like urticaria. Metronidazole is the standard first-line treatment; however, the emergence of resistance and instances of treatment failure have prompted research into alternative antiparasitic therapies. Understanding the pathogenic potential of Blastocystis and optimizing treatment strategies are essential for managing this prevalent infection.

| Associated when high | | | |
|---|---|---|---|
| *g. Prevotella (f. Paraprevotellaceae)* | Very High | 0 15.471 16.709 | Unknown |

Out of 16 possible associations, 4 were detected in the patient; among these, 1 is outside the normal range.

#### Critical Illness

Critical illness refers to a state of severe physiological compromise requiring intensive medical intervention, often characterized by multi-organ dysfunction and high mortality risk. Its clinical significance lies in the urgent need for effective management strategies to improve patient outcomes, such as nutrition protocols in intensive care settings.

| Associated when high | | | |
|---|---|---|---|
| *p. Proteobacteria* | Very High | 0 11.493 12.413 | Commensal |

| Associated when low | | | |
|---|---|---|---|
| *g. Blautia* | Very Low | 0 0 30.776 | Commensal |

Out of 9 possible associations, 3 were detected in the patient; among these, 2 are outside the normal range.

#### Hospital Admission; Anorexia Nervosa

+55 84 3113 5085

www.dnagtxbioinfo.com

Anorexia nervosa (AN) is a severe psychiatric disorder marked by extreme caloric restriction and significant weight loss, leading to detrimental physiological changes, particularly in skeletal muscle. Its clinical significance lies in the persistent alterations in muscle proteomics and function, even after weight restoration, highlighting the complexity of recovery and the need for targeted interventions.

| Associated when high | | | |
|---|---|---|---|
| *g. Lachnospira* | Very High | 0 — 5.328 — 5.754 | Commensal |

#### Kidney Calculi

Kidney calculi, or kidney stones, are solid mineral deposits that form in the renal system, often leading to pain, urinary obstruction, and potential complications such as recurrent infections or kidney damage. Their clinical significance lies in their association with conditions like urinary tract infections and the potential need for surgical intervention, as seen in cases involving kidney transplants.

| Associated when high | | | |
|---|---|---|---|
| *g. Sutterella* | Very High | 0 — 12.441 — 13.437 | Unknown |

Out of 18 possible associations, 2 were detected in the patient; among these, 1 is outside the normal range.

#### Non-Edematous Severe Acute Malnutrition

Non-edematous refers to a condition characterized by weight gain without the presence of fluid retention or swelling in tissues. Clinically, it indicates a true increase in body mass, often linked to metabolic changes, particularly in the context of recovery from malnutrition.

| Associated when low | | | |
|---|---|---|---|
| *f. Lachnospiraceae* | Very Low | 0 — 6.439 — 67.829 | Unknown |



BRAZIL | UAE | USA
Av. Senador Salgado Filho, 3.000 - Room A402 – CEP: 59.078-970 – Natal / RN
Av. Paulista, 1.636 - Conjunto 301 – CEP: 13.102-000 – São Paulo / SP
DNA GTx

+55 84 3113 5085

www.dnagtxbioinfo.com

### 5. Abundance of Identified Bacteria

#### Pathogenic Bacteria

| Bacteria | Percentile | Percentage | Level |
|---|---|---|---|
| *o. Burkholderiales* | 99.72 | 11.4934% | Very High |

| Bacteria | Percentile | Percentage | Level |
|---|---|---|---|

#### Commensal Bacteria

| Bacteria | Percentile | Percentage | Level |
|---|---|---|---|
| *c. Bacteroidia* | 99.72 | 56.6458% | Very High |
| *o. Bacteroidales* | 99.72 | 56.6458% | Very High |
| *f. Bacteroidaceae* | 99.72 | 39.0774% | Very High |
| *g. Bacteroides (f. Bacteroidaceae)* | 99.72 | 42.3004% | Very High |
| *Bacteroides plebeius* | 99.72 | 67.1261% | Very High |
| *g. Odoribacter* | 99.43 | 0.7871% | Very High |
| *Parabacteroides distasonis* | 93.73 | 2.5708% | Very High |
| *Alistipes indistinctus* | 99.43 | 1.641% | Very High |

+55 84 3113 5085

www.dnagtxbioinfo.com

| | | | |
|---|---|---|---|
| *c. Clostridia* | 0.00 | 29.4605% | Very Low |
| *g. Blautia* | 0.00 | 0% | Very Low |
| *g. Lachnospira* | 98.29 | 5.3277% | Very High |
| *g. Gemmiger* | 98.29 | 7.5579% | Very High |
| *Gemmiger formicilis* | 99.43 | 19.5383% | Very High |
| *g. Oscillospira* | 96.01 | 9.9319% | Very High |

| Bacteria | Percentile | Percentage | Level |
|---|---|---|---|
| *Bacteroides uniformis* | 10.54 | 0.186% | Normal |
| *g. Parabacteroides* | 87.75 | 1.8027% | Normal |
| *f. Prevotellaceae* | 57.26 | 0.2971% | Normal |
| *g. Prevotella (f. Prevotellaceae)* | 58.40 | 0.3216% | Normal |
| *Prevotella copri* | 70.66 | 0.8314% | Normal |
| *f. Rikenellaceae* | 55.56 | 0.5864% | Normal |
| *g. Alistipes* | 68.95 | 0.6348% | Normal |
| *g. Coprococcus* | 0.57 | 0.0042% | Normal |
| *f. Ruminococcaceae* | 24.79 | 23.0219% | Normal |
| *g. Butyricicoccus* | 31.62 | 0.5374% | Normal |
| *Butyricicoccus pullicaecorum* | 48.15 | 1.3893% | Normal |
| *c. Erysipelotrichi* | 38.18 | 2.4003% | Normal |
| *f. Erysipelotrichaceae* | 36.75 | 2.4003% | Normal |
| *g. Eubacterium* | 82.62 | 2.5983% | Normal |
| *Eubacterium biforme* | 86.89 | 6.717% | Normal |

#### Bacteria With Uncertain Effects

| Bacteria | Percentile | Percentage | Level |
|---|---|---|---|

+55 84 3113 5085

www.dnagtxbioinfo.com

| | | | |
|---|---|---|---|
| *f. Odoribacteraceae* | 98.86 | 0.7271% | Very High |
| *f. Paraprevotellaceae* | 99.72 | 14.2924% | Very High |
| *g. Prevotella (f. Paraprevotellaceae)* | 99.72 | 15.4712% | Very High |
| *o. Clostridiales* | 0.00 | 29.4605% | Very Low |
| *f. Lachnospiraceae* | 0.00 | 6.4386% | Very Low |
| *c. Betaproteobacteria* | 99.72 | 11.4934% | Very High |
| *f. Alcaligenaceae* | 99.72 | 11.4934% | Very High |
| *g. Sutterella* | 99.72 | 12.4413% | Very High |

| Bacteria | Percentile | Percentage | Level |
|---|---|---|---|
| *f. Porphyromonadaceae* | 89.74 | 1.6654% | Normal |
| *g. Ruminococcus (f. Ruminococcaceae)* | 4.56 | 0.2835% | Normal |
| *o. Erysipelotrichales* | 38.18 | 2.4003% | Normal |

#### 6. References:

[1] Mucosal adherent bacterial dysbiosis in patients with colorectal adenomas (PMID:27194068);

[2] The Intestinal Microbiota in Acute Anorexia Nervosa and During Renourishment: Relationship to Depression, Anxiety, and Eating Disorder Psychopathology (PMID:26428446);

[3] Human colonic microbiota associated with diet, obesity and weight loss. (10.1038/ijo.2008.155);

+55 84 3113 5085

www.dnagtxbioinfo.com

[4] Arabinogalactan and fructo-oligosaccharides have a different fermentation profile in the Simulator of the Human Intestinal Microbial Ecosystem (SHIME). (10.1111/1758-2229.12056);

[5] Resveratrol alleviates temporomandibular joint inflammatory pain by recovering disturbed gut microbiota. (10.1016/j.bbi.2020.01.016);

[6] The Defect in Regulatory T Cells in Psoriasis and Therapeutic Approaches. (10.3390/jcm10173880);

[7] Dietary Probiotics or Synbiotics Supplementation During Gestation, Lactation, and Nursery Periods Modifies Colonic Microbiota, Antioxidant Capacity, and Immune Function in Weaned Piglets. (10.3389/fvet.2020.597832);

[8] Gut microbiome in ADHD and its relation to neural reward anticipation (PMID:28863139);

[9] Insight into alteration of gut microbiota in Clostridium difficile infection and asymptomatic C. difficile colonization (PMID:25817005);

[10] Gut microbiota trajectory in early life may predict development of celiac disease (PMID:29458413);

[11] Low counts of Faecalibacterium prausnitzii in colitis microbiota (PMID:19235886);

[12] Alterations in the gut microbiome of children with severe ulcerative colitis (PMID:22170749);

[13] Mucosa-associated microbiota signature in colorectal cancer (PMID:28600626);

[14] Increased rectal microbial richness is associated with the presence of colorectal adenomas in humans (PMID:22622349);

[15] Gut microbiota in human adults with type 2 diabetes differs from non-diabetic adults (PMID:20140211);

[16] The stool microbiota of insulin resistant women with recent gestational diabetes, a high risk group for type 2 diabetes (PMID:26279179);

[17] 16S rRNA gene-based analysis of fecal microbiota from preterm infants with and without necrotizing enterocolitis (PMID:19369970);

[18] Intestinal dysbiosis associated with systemic lupus erythematosus (PMID:25271284);

[19] Intestinal dysbiosis in preterm infants preceding necrotizing enterocolitis: a systematic review and meta-analysis (PMID:28274256);

[20] Maternal antenatal treatments influence initial oral microbial acquisition in preterm infants. (10.1055/s-0032-1321499);

[21] Characterization of the cellulolytic bacteria communities along the gastrointestinal tract of Chinese Mongolian sheep by using PCR-DGGE and real-time PCR analysis. (10.1007/s11274-015-1860-z);

[22] Dietary xylo-oligosaccharide supplementation alters gut microbial composition and activity in pigs according to age and dose. (10.1186/s13568-019-0858-6);

[23] Seaweed polysaccharides treatment alleviates injury of inflammatory responses and gut barrier in LPS-induced mice. (10.1016/j.micpath.2023.106159);

[24] Disruption of the human gut microbiota following Norovirus infection (PMID:23118957);

[25] Gut microbiota dysbiosis and bacterial community assembly associated with cholesterol gallstones in large-scale study (PMID:24083370);

[26] Fecal microbial dysbiosis in Chinese patients with inflammatory bowel disease (PMID:29632427);

[27] Critically ill patients demonstrate large interpersonal variation in intestinal microbiota dysregulation: a pilot study (PMID:27837233);

[28] Ketogenic diet poses a significant effect on imbalanced gut microbiota in infants with refractory epilepsy (PMID:28970732);

[29] Associations of cocaine use and HIV infection with the intestinal microbiota, microbial translocation, and inflammation (PMID:24650829);

[30] Dysbiosis Signatures of Gut Microbiota Along the Sequence from Healthy, Young Patients to Those with Overweight and Obesity (PMID:29280312);

[31] Alterations in the gut microbiome of children with severe ulcerative colitis. (10.1002/ibd.22860);

[32] Insight into alteration of gut microbiota in Clostridium difficile infection and asymptomatic C. difficile colonization. (10.1016/j.anaerobe.2015.03.008);

[33] Sodium butyrate ameliorates thiram-induced tibial dyschondroplasia and gut microbial dysbiosis in broiler chickens. (10.1016/j.ecoenv.2022.114134);

[34] Effect of Extracelluar Vesicles Derived from Lactobacillus plantarum Q7 on Gut Microbiota and Ulcerative Colitis in Mice. (10.3389/fimmu.2021.777147);

[35] Cellular aspects of Burkholderia cepacia infection. (10.1016/s1286-4579(01)01389-2);

[36] The multifarious, multireplicon Burkholderia cepacia complex. (10.1038/nrmicro1085);

[37] Antibiotic resistance in Burkholderia species. (10.1016/j.drup.2016.07.003);

[38] Burkholderia pseudomallei. (10.1016/j.tim.2023.07.008);

[39] Gut microbiome of Moroccan colorectal cancer patients. (10.1007/s00430-018-0542-5);

[40] Uremic Toxin-Producing Gut Microbiota in Rats with Chronic Kidney Disease. (10.1159/000450619);

[41] Efficacy of intestinal microorganisms on immunotherapy of non-small cell lung cancer. (10.1016/j.heliyon.2024.e29899);

[42] He who controls Clostridia and Bacteroidia controls the gut microbiome: The concept of targeted probiotics to restore the balance of keystone taxa in irritable bowel syndrome. (10.1111/nmo.14805);

[43] Correlation between the human fecal microbiota and depression (PMID:24888394);

[44] Sodium Butyrate Ameliorates Gut Microbiota Dysbiosis in Lupus-Like Mice. (10.3389/fnut.2020.604283);

[45] Gut microbiota-based discriminative model for patients with ulcerative colitis: A meta-analysis and real-world study.

(10.1097/MD.0000000000037091);

[**46**] Analysis of gut microbiome composition, function, and phenotype in patients with osteoarthritis. (10.3389/fmicb.2022.980591);

[**47**] Rett Syndrome: A Focus on Gut Microbiota (PMID:28178201);

[**48**] Anticancer immunotherapy by CTLA-4 blockade relies on the gut microbiota. (10.1126/science.aad1329);

[**49**] The evolution of cooperation within the gut microbiota. (10.1038/nature17626);

[**50**] Structural modulation of gut microbiota by chondroitin sulfate and its oligosaccharide. (10.1016/j.ijbiomac.2016.04.091);

[**51**] Cross-feeding between intestinal pathobionts promotes their overgrowth during undernutrition. (10.1038/s41467-021-27191-x);

[**52**] Streamlined Genetic Manipulation of Diverse Bacteroides and Parabacteroides Isolates from the Human Gut Microbiota. (10.1128/mBio.01762-19);

[**53**] Gut microbiota composition and Clostridium difficile infection in hospitalized elderly individuals: a metagenomic study (PMID:27166072);

[**54**] Microbiome data distinguish patients with Clostridium difficile infection and non-C. difficile-associated diarrhea from healthy controls (PMID:24803517);

[**55**] Alterations of mucosal microbiota in the colon of patients with inflammatory bowel disease revealed by real time polymerase chain reaction amplification of 16S ribosomal ribonucleic acid (PMID:26261163);

[**56**] Tumour-associated and non-tumour-associated microbiota in colorectal cancer (PMID:26992426);

[**57**] Comparison of the fecal microbiota profiles between ulcerative colitis and Crohn’s disease using terminal restriction fragment length polymorphism analysis (PMID:21253779);

[**58**] Fecal microbiota imbalance in Mexican children with type 1 diabetes (PMID:24448554);

[**59**] Gut microbiota profiling in Han Chinese with type 1 diabetes (PMID:29733871);

[**60**] Mucosa-associated bacterial diversity in necrotizing enterocolitis (PMID:25203729);

[**61**] Integrated analysis of microbiome and host transcriptome reveals correlations between gut microbiota and clinical outcomes in HBV-related hepatocellular carcinoma (PMID:33225985);

[**62**] Association study of gut flora in Wilson's disease through high-throughput sequencing (PMID:30075590);

[**63**] Leveraging sequence-based faecal microbial community survey data to identify a composite biomarker for colorectal cancer (PMID:28341746);

[**64**] Gut Microbiota Serves a Predictable Outcome of Short-Term Low-Carbohydrate Diet (LCD) Intervention for Patients with Obesity. (10.1128/Spectrum.00223-21);

[**65**] Ligustrum robustum Intake, Weight Loss, and Gut Microbiota: An Intervention Trial. (10.1155/2019/4643074);

[**66**] Betaine inhibits Toll-like receptor 4 responses and restores intestinal microbiota in acute liver failure mice. (10.1038/s41598-020-78935-6);

[**67**] California strawberry consumption increased the abundance of gut microorganisms related to lean body weight, health and longevity in healthy subjects. (10.1016/j.nutres.2020.12.006);

[**68**] Gut Microbiota Markers and Dietary Habits Associated with Extreme Longevity in Healthy Sardinian Centenarians. (10.3390/nu14122436);

[**69**] Orthogonal Dietary Niche Enables Reversible Engraftment of a Gut Bacterial Commensal. (10.1016/j.celrep.2018.07.032);

[**70**] Gut Microbiota Analysis in Postoperative Lynch Syndrome Patients. (10.3389/fmicb.2019.01746);

[**71**] Alterations of the Gut Microbiota in Multiple System Atrophy Patients. (10.3389/fnins.2019.01102);

[**72**] US nativity and dietary acculturation impact the gut microbiome in a diverse US population. (10.1016/j.s41396-020-0630-6);

[**73**] Gut metagenomic and short chain fatty acids signature in hypertension: a cross-sectional study. (10.1038/s41598-020-63475-w);

[**74**] Characterization of BpGH16A of Bacteroides plebeius, a key enzyme initiating the depolymerization of agarose in the human gut. (10.1007/s00253-020-11039-3);

[**75**] Gut microbiome profile in psoriatic patients treated and untreated with biologic therapy. (10.1111/1346-8138.15680);

[**76**] Bacteroides plebeius improves muscle wasting in chronic kidney disease by modulating the gut-renal muscle axis. (10.1111/jcmm.17626);

[**77**] Multi-cohort analysis of depression-associated gut bacteria sheds insight on bacterial biomarkers across populations. (10.1007/s00018-022-04650-2);

[**78**] Gut colonization of Bacteroides plebeius suppresses colitis-associated colon cancer development. (10.1128/spectrum.02599-24);

[**79**] Characteristics of Faecal Microbiota in Paediatric Crohn’s Disease and Their Dynamic Changes During Infliximab Therapy (PMID:29194468);

[**80**] Characteristics of fecal microbiota in non-alcoholic fatty liver disease patients (PMID:29948900);

[**81**] [Role of gut microbiota in children with allergic rhinitis with high serum total IgE level]. (10.13201/j.issn.2096-7993.2020.12.016);

[**82**] Gut Microbiome Is Related to Cognitive Impairment in Peritoneal Dialysis Patients. (10.3390/nu16162659);

[**83**] Characterization of the blood microbiota in children with Celiac disease. (10.1016/j.crmicr.2021.100069);

[**84**] Gut bacteria from multiple sclerosis patients modulate human T cells and exacerbate symptoms in mouse models. (10.1073/pnas.1711235114);

[**85**] Parabacteroides distasonis Alleviates Obesity and Metabolic Dysfunctions via Production of Succinate and Secondary Bile Acids.

+55 84 3113 5085

www.dnagtxbioinfo.com

(10.1016/j.celrep.2018.12.028);

[**86**] Roles of intestinal Parabacteroides in human health and diseases. (10.1093/femsle/fnac072);

[**87**] Potential roles of gut microbiome and metabolites in modulating ALS in mice. (10.1038/s41586-019-1443-5);

[**88**] Gut Microbiota and Gestational Diabetes Mellitus: A Review of Host-Gut Microbiota Interactions and Their Therapeutic Potential. (10.3389/fcimb.2020.00188);

[**89**] Administration of Alistipes indistinctus prevented the progression from nonalcoholic fatty liver disease to nonalcoholic steatohepatitis by enhancing the gut barrier and increasing Lactobacillus spp. (10.1016/j.bbrc.2024.151033);

[**90**] Alistipes indistinctus-derived hippuric acid promotes intestinal urate excretion to alleviate hyperuricemia. (10.1016/j.chom.2024.02.001);

[**91**] The Genus Alistipes: Gut Bacteria With Emerging Implications to Inflammation, Cancer, and Mental Health. (10.3389/fimmu.2020.00906);

[**92**] Advanced glycation end products dietary restriction effects on bacterial gut microbiota in peritoneal dialysis patients; a randomized open label controlled trial. (10.1371/journal.pone.0184789);

[**93**] Human gut microbiome and risk for colorectal cancer (PMID:24316595);

[**94**] Gut bacteria dysbiosis and necrotising enterocolitis in very low birthweight infants: a prospective case-control study (PMID:26969089);

[**95**] Human gut microbiota in obesity and after gastric bypass (PMID:19164560);

[**96**] Human gut microbiome and risk for colorectal cancer. (10.1093/jnci/djt300);

[**97**] Consumption of lysozyme-rich milk can alter microbial fecal populations. (10.1128/AEM.00956-12);

[**98**] Association of fecal microbial diversity and taxonomy with selected enzymatic functions. (10.1371/journal.pone.0039745);

[**99**] Patients with Acne Vulgaris Have a Distinct Gut Microbiota in Comparison with Healthy Controls (PMID:29756631);

[**100**] Gut Microbiota Composition and Fecal Metabolic Phenotype in Patients With Acute Anterior Uveitis (PMID:29625474);

[**101**] Gut microbiota dysbiosis in depressed women: The association of symptom severity and microbiota function (PMID:33421868);

[**102**] Structural changes of gut microbiota in Parkinson's disease and its correlation with clinical features (PMID:28536926);

[**103**] Structural changes of gut microbiota in Parkinson's disease and its correlation with clinical features. (10.1007/s11427-016-9001-4);

[**104**] Immunity improvement and gut microbiota remodeling of mice by wheat germ globulin. (10.1007/s11274-021-03034-1);

[**105**] Gut dysbiosis is associated with the reduced exercise capacity of elderly patients with hypertension. (10.1038/s41440-018-0110-9);

[**106**] Protective effects of Tibetan kefir in mice with ochratoxin A-induced cecal injury. (10.1016/j.foodres.2022.111551);

[**107**] Identification of Gut Microbiota and Metabolites Signature in Patients With Irritable Bowel Syndrome (PMID:31681624);

[**108**] Phenylketonuria and Gut Microbiota: A Controlled Study Based on Next-Generation Sequencing. (10.1371/journal.pone.0157513);

[**109**] Alterations in the gut bacterial microbiome in fungal Keratitis patients. (10.1371/journal.pone.0199640);

[**110**] Dysbiosis in the Gut Bacterial Microbiome of Patients with Uveitis, an Inflammatory Disease of the Eye. (10.1007/s12088-018-0746-9);

[**111**] Fecal Microbiomes Distinguish Patients With Autoimmune Hepatitis From Healthy Individuals. (10.3389/fcimb.2020.00342);

[**112**] Clinical Phenotypes of Parkinson's Disease Associate with Distinct Gut Microbiota and Metabolome Enterotypes. (10.3390/biom11020144);

[**113**] Lachnospira is a signature of antihistamine efficacy in chronic spontaneous urticaria. (10.1111/exd.14460);

[**114**] Altered Gut Microbiota Composition Associated with Eczema in Infants (PMID:27812181);

[**115**] Predictive Metagenomic Analysis of Autoimmune Disease Identifies Robust Autoimmunity and Disease Specific Microbial Signatures. (10.3389/fmicb.2021.621310);

[**116**] Postoperative Changes in Fecal Bacterial Communities and Fermentation Products in Obese Patients Undergoing Bilio-Intestinal Bypass. (10.3389/fmicb.2016.00200);

[**117**] Gut Microbiota Dysbiosis in Patients with Endometrial Cancer vs. Healthy Controls Based on 16S rRNA Gene Sequencing. (10.1007/s00284-023-03361-6);

[**118**] Microbial Community of Healthy Thai Vegetarians and Non-Vegetarians, Their Core Gut Microbiota, and Pathogen Risk. (10.4014/jmb.1603.03057);

[**119**] Dietary Taxifolin Protects Against Dextran Sulfate Sodium-Induced Colitis via NF-B Signaling, Enhancing Intestinal Barrier and Modulating Gut Microbiota. (10.3389/fimmu.2020.631809);

[**120**] Influence of Short-Term Consumption of Hericium erinaceus on Serum Biochemical Markers and the Changes of the Gut Microbiota: A Pilot Study. (10.3390/nu13031008);

[**121**] Vitamin C Supplementation in Healthy Individuals Leads to Shifts of Bacterial Populations in the Gut-A Pilot Study. (10.3390/antiox10081278);

[**122**] Self-Initiated Dietary Adjustments Alter Microbiota Abundances: Implications for Perceived Health. (10.3390/nu16203544);

[**123**] Gut microbiome of Moroccan colorectal cancer patients (PMID:29687353);

[**124**] Analysis of Gut Microbiota in Patients with Parkinson's Disease (PMID:28429209);

[**125**] Lactobacillus rhamnosus GG-supplemented formula expands butyrate-producing bacterial strains in food allergic infants. (10.1038/ismej.2015.151);

**[126]** Gut Microbiota Co-microevolution with Selection for Host Humoral Immunity. (10.3389/fmicb.2017.01243);

**[127]** The association between gut microbiome and anthropometric measurements in Bangladesh. (10.1080/19490976.2019.1614394);

**[128]** Gut Microbiota and Risk of Persistent Nonalcoholic Fatty Liver Diseases. (10.3390/jcm8081089);

**[129]** Korean Traditional Medicine (Jakyakgamcho-tang) Ameliorates Colitis by Regulating Gut Microbiota. (10.3390/metabo9100226);

**[130]** Rectal microbiota among HIV-uninfected, untreated HIV, and treated HIV-infected in Nigeria (PMID:28118207);

**[131]** Reduced microbiome alpha diversity in young patients with ADHD (PMID:30001426);

**[132]** Colonization with the enteric protozoa Blastocystis is associated with increased diversity of human gut bacterial microbiota (PMID:27147260);

**[133]** Molecular profiling of mucosal tissue associated microbiota in patients manifesting acute exacerbations and remission stage of ulcerative colitis (PMID:29796862);

**[134]** Smokers with Active Crohn's Disease Have a Clinically Relevant Dysbiosis of the Gastrointestinal Microbiota (PMID:22102318);

**[135]** Prevotella and Klebsiella proportions in fecal microbial communities are potential characteristic parameters for patients with major depressive disorder (PMID:27741466);

**[136]** Obesity Alters the Microbial Community Profile in Korean Adolescents (PMID:26230509);

**[137]** Phenylketonuria and Gut Microbiota: A Controlled Study Based on Next-Generation Sequencing (PMID:27336782);

**[138]** The Intratumor Microbiota Signatures Associate With Subtype, Tumor Stage, and Survival Status of Esophageal Carcinoma. (10.3389/fonc.2021.754788);

**[139]** Alterations in the gut microbiome and metabolism with coronary artery disease severity (PMID:31027508);

**[140]** Restitution of gut microbiota in Ugandan children administered with probiotics (Lactobacillus rhamnosus GG and Bifidobacterium animalis subsp. lactis BB-12) during treatment for severe acute malnutrition (PMID:31959047);

**[141]** The distinct features of microbial 'dysbiosis' of Crohn's disease do not occur to the same extent in their unaffected, genetically-linked kindred (PMID:28222161);

**[142]** Gut microbiota in early pediatric multiple sclerosis: a case-control study (PMID:27176462);

**[143]** Legume-nodulating betaproteobacteria: diversity, host range, and future prospects. (10.1094/MPMI-06-11-0172);

**[144]** Genome implosion elicits host-confinement in Alcaligenaceae: evidence from the comparative genomics of Tetrathiobacter kashmirensis, a pathogen in the making. (10.1371/journal.pone.0064856);

**[145]** Resemblance and divergence: the "new" members of the genus Bordetella. (10.1007/s00430-010-0148-z);

**[146]** Virulence factor secretion and translocation by Bordetella species. (10.1016/j.mib.2009.01.001);

**[147]** Bordetella holmesii: Still Emerging and Elusive 20 Years On. (10.1128/microbiolspec.EI10-0003-2015);

**[148]** Assessing gut microbiota perturbations during the early phase of infectious diarrhea in Vietnamese children (PMID:28767339);

**[149]** 16S rRNA gene sequencing reveals altered composition of gut microbiota in individuals with kidney stones (PMID:29353409);

**[150]** Human Gut Microbiota Associated with Obesity in Chinese Children and Adolescents (PMID:29214176);

**[151]** Application of novel PCR-based methods for detection, quantitation, and phylogenetic characterization of Sutterella species in intestinal biopsy samples from children with autism and gastrointestinal disturbances. (10.1128/mBio.00261-11);

**[152]** The genus Sutterella is a potential contributor to glucose metabolism improvement after Roux-en-Y gastric bypass surgery in T2D. (10.1016/j.diabres.2020.108116);

**[153]** Structural Alteration of Gut Microbiota during the Amelioration of Human Type 2 Diabetes with Hyperlipidemia by Metformin and a Traditional Chinese Herbal Formula: a Multicenter, Randomized, Open Label Clinical Trial (PMID:29789365);

**[154]** Impact of enrofloxacin on the human intestinal microbiota revealed by comparative molecular analysis (PMID:22321759);

**[155]** Obesity and mental health improvement following nutritional education focusing on gut microbiota composition in Japanese women: a randomised controlled trial (PMID:30523432);

**[156]** The Effect of Probiotics on Gut Microbiota during the Helicobacter pylori Eradication: Randomized Controlled Trial (PMID:26395781);

**[157]** Gut microbiota is associated with adiposity markers and probiotics may impact specific genera (PMID:31250099);

**[158]** Differential Changes in Gut Microbiota After Gastric Bypass and Sleeve Gastrectomy Bariatric Surgery Vary According to Diabetes Remission (PMID:27738970);

**[159]** Flavonoid-Rich Orange Juice Intake and Altered Gut Microbiome in Young Adults with Depressive Symptom: A Randomized Controlled Study (PMID:32570775);

**[160]** Rifaximin alters gut microbiota profile, but does not affect systemic inflammation - a randomized controlled trial in common variable immunodeficiency (PMID:30655568);

**[161]** Gut microbiota composition in relation to the metabolic response to 12-week combined polyphenol supplementation in overweight men and women (PMID:28589947);

**[162]** Long Term Development of Gut Microbiota Composition in Atopic Children: Impact of Probiotics (PMID:26378926);

**[163]** Deep T ranscranial Magnetic Stimulation Affects Gut Microbiota Composition in Obesity: Results of Randomized Clinical T rial (PMID:33946648);

+55 84 3113 5085

www.dnagtxbioinfo.com

[164] Evaluation of the effects of intrapartum antibiotic prophylaxis on newborn intestinal microbiota using a sequencing approach targeted to multi hypervariable 16S rDNA regions (PMID:26971496);

[165] Impact of intrapartum antimicrobial prophylaxis upon the intestinal microbiota and the prevalence of antibiotic resistance genes in vaginally delivered full-term neonates (PMID:28789705);

[166] Probiotics modify human intestinal mucosa-associated microbiota in patients with colorectal cancer (PMID:26238090);

[167] Effect of Synbiotic Supplementation in a Very-Low-Calorie Ketogenic Diet on Weight Loss Achievement and Gut Microbiota: A Randomized Controlled Pilot Study (PMID:31298466);

[168] 454 Pyrosequencing Reveals a Shift in Fecal Microbiota of Healthy Adult Men Consuming Polydextrose or Soluble Corn Fiber (PMID:22649263);

[169] The altered gut microbiota in adults with cystic fibrosis (PMID:28279152);

[170] Alterations of gut microbiota composition in post-finasteride patients: a pilot study (PMID:32951160);

[171] Iron in Micronutrient Powder Promotes an Unfavorable Gut Microbiota in Kenyan Infants (PMID:28753958);

[172] The therapeutic efficacy of Bifidobacterium animalis subsp. lactis BB-12 ® in infant colic: A randomised, double blind, placebo-controlled trial (PMID:31797399);

[173] Dietary intake of fat and fibre according to reference values relates to higher gut microbiota richness in overweight pregnant women (PMID:28901891);

[174] Fermented Milk Containing Lactobacillus casei Strain Shirota Preserves the Diversity of the Gut Microbiota and Relieves Abdominal Dysfunction in Healthy Medical Students Exposed to Academic Stress (PMID:27208120);

[175] Helicobacter pylori eradication with bismuth quadruple therapy leads to dysbiosis of gut microbiota with an increased relative abundance of Proteobacteria and decreased relative abundances of Bacteroidetes and Actinobacteria (PMID:29897654);

[176] Gut microbiota associations with diet in irritable bowel syndrome and the effect of low FODMAP diet and probiotics (PMID:33183883);

[177] Metabolic phenotypes and the gut microbiota in response to dietary resistant starch type 2 in normalweight subjects: a randomized crossover trial (PMID:30894560);

[178] The effect of drinking water pH on the human gut microbiota and glucose regulation: results of a randomized controlled cross-over intervention (PMID:30413727);

[179] Gut microbiota composition after diet and probiotics in overweight breast cancer survivors: a randomized open-label pilot intervention trial (PMID:32234652);

[180] Treatment-Specific Composition of the Gut Microbiota Is Associated With Disease Remission in a Pediatric Crohn's Disease Cohort (PMID:31276165);

[181] Arabinoxylan oligosaccharides and polyunsaturated fatty acid effects on gut microbiota and metabolic markers in overweight individuals with signs of metabolic syndrome: A randomized cross-over trial (PMID:30827722);

[182] Effects of probiotic supplementation on serum trimethylamine-N-oxide level and gut microbiota composition in young males: a double-blinded randomized controlled trial (PMID:32440731);

[183] Probiotic supplementation restores normal microbiota composition and function in antibiotic-treated and in caesarean-born infants (PMID:30326954);

[184] Influence of a 3-month low-calorie Mediterranean diet compared to the vegetarian diet on human gut microbiota and SCFA: the CARDIVEG Study (PMID:31292752);

[185] Gut Microbial Diversity in Antibiotic-Naive Children After Systemic Antibiotic Exposure: A Randomized Controlled Trial (PMID:28402408);

[186] Effects of dietary fat on gut microbiota and faecal metabolites, and their relationship with cardiometabolic risk factors: a 6-month randomised controlled-feeding trial (PMID:30782617);

[187] A randomised trial of the effect of omega-3 polyunsaturated fatty acid supplements on the human intestinal microbiota (PMID:28951525);

[188] Impact of probiotics supplement on the gut microbiota in neonates with antibiotic exposure: an open-label single-center randomized parallel controlled study (PMID:34331676);

[189] Elucidating the gut microbiome of ulcerative colitis: bifidobacteria as novel microbial biomarkers (PMID:27604252);

[190] Probiotic or synbiotic alters the gut microbiota and metabolism in a randomised controlled trial of weight management in overweight adults (PMID:30525950);

[191] Impact of dietary fiber supplementation on modulating microbiota–host–metabolic axes in obesity (PMID:30572270);

[192] Effects of a Vegetarian Diet on Cardiometabolic Risk Factors, Gut Microbiota, and Plasma Metabolome in Subjects With Ischemic Heart Disease: A Randomized, Crossover Study (PMID:32893710);

[193] Differences in gut microbiota profile between women with active lifestyle and sedentary women (PMID:28187199);

[194] Differential effects of antiretrovirals on microbial translocation and gut microbiota composition of HIV-infected patients (PMID:28362071);

[195] Effects of Synbiotic Supplement on Human Gut Microbiota, Body Composition and Weight Loss in Obesity (PMID:31952249);

[196] Effects of fermented soymilk with Lactobacillus casei Shirota on skin condition and the gut microbiota: a randomised clinical pilot trial (PMID:29264969);

[197] Randomized controlled trial on the impact of early-life intervention with bifidobacteria on the healthy infant fecal microbiota and metabolome (PMID:28877893);


Av. Senador Salgado Filho, 3.000 - Room A402 – CEP: 59.078-970 – Natal / RN
Av. Paulista, 1.636 - Conjunto 301 – CEP: 13.102-000 – São Paulo / SP

+55 84 3113 5085

www.dnagtxbioinfo.com

[**198**] Impact of probiotic Saccharomyces boulardii on the gut microbiome composition in HIV-treated patients: A double-blind, randomised, placebo-controlled trial (PMID:28388647);

[**199**] Soluble Corn Fiber Increases Calcium Absorption Associated with Shifts in the Gut Microbiome: A Randomized Dose-Response Trial in Free-Living Pubertal Females (PMID:27281813);

[**200**] A Vegetarian Diet Is a Major Determinant of Gut Microbiota Composition in Early Pregnancy (PMID:30002323);

[**201**] Synbiotics Easing Renal Failure by Improving Gut Microbiology (SYNERGY): A Randomized Trial (PMID:26772193);
